# Integrated Omics Analyses Reveal Differential Gene Expression and Potential for Cooperation Between Denitrifying Polyphosphate and Glycogen Accumulating Organisms

**DOI:** 10.1101/2020.01.10.901413

**Authors:** Yubo Wang, Han Gao, George Wells

## Abstract

Unusually high accumulation of the potent greenhouse gas nitrous oxide (N_2_O) has previously been documented in denitrifying biological phosphorus (P) removal bioprocesses, but the roles of differential denitrification gene expression patterns and ecological interactions between key functional groups in driving these emissions are not well understood. To address these knowledge gaps, we applied genome-resolved metagenomics and metatranscriptomics to a denitrifying bioprocess enriched in as-yet-uncultivated denitrifying polyphosphate accumulating organisms (PAOs) affiliated with *Candidatus* Accumulibacter. The 6 transcriptionally most active populations in the community included three co-occurring Accumulibacter strains affiliated with clades IF (a novel clade identified in this study), IA, and IC, and a competing glycogen accumulating organism (GAO) affiliated with *Candidatus* Competibacter. Strongly elevated expression of nitrite reductase compared to nitrous oxide reductase was observed in the overall community and in Accumulibacter populations, suggesting a strong role for differential gene expression in driving N_2_O accumulation. Surprisingly, while ∼90% of nitrite reductase gene transcripts mapped to the three co-occurring PAO populations, ∼93% of nitric oxide reductase gene transcripts were expressed by the GAO population. This suggests the potential for cooperation between GAOs and PAOs in reducing denitrification intermediates. Such cooperation may benefit the community by reducing the accumulation of toxic nitric oxide.

**Originality-Significance Statement:** Polyphosphate accumulating organisms (PAOs) affiliated with as-yet-uncultivated *Ca.* ‘Accumulibacter phosphatis’ are increasingly employed in enhanced biological phosphorus removal (EBPR) processes, a common environmental biotechnology for removing phosphorus from wastewater and thereby preventing detrimental impacts of nutrient pollution. Under anoxic conditions, PAOs have been associated with unusually high emissions of the potent greenhouse gas and denitrification intermediate nitrous oxide. However, the underlying mechanisms and biological controls on incomplete denitrification by denitrifying Accumulibacter, their ecological interactions with understudied glycogen accumulating organisms (GAOs), and patterns of gene expression under anoxic conditions are all poorly understood. Here, we describe genomic features of a previously unrecognized clade of Accumulibacter that is putatively adapted to high rate P uptake under nitrite-driven denitrification and provide evidence that differential gene expression (namely elevated expression of nitrite reductase compared to nitrous oxide reductase) by Accumulibacter is a key control on nitrous oxide production. Moreover, we document genomic and transcriptional potential for cooperation and crossfeeding of the denitrification intermediate nitric oxide between GAOs and PAOs. This is surprising because GAOs are conventionally considered to be competitors to PAOs, and because nitric oxide is toxic to most microorganisms at low concentrations. Taken together, our work provides significant new understanding of metabolic and ecological interactions in EBPR processes that are critical to environmental protection; demonstrates the potential of previously unrecognized crossfeeding of the denitrification intermediate nitric oxide; and expands our understanding of genomic features and clade level diversity of Accumulibacter.

## Introduction

Enhanced biological phosphorus removal (EBPR) bioprocesses that rely on as-yet-uncultivated polyphosphate accumulating organisms (PAOs) are increasingly used for sustainable phosphorus (P) removal and recovery from wastewater (Stokholm-Bjerregaard et al., 2017). The most common PAO in most full- and lab-scale EBPR processes affiliates with *Candidatus* ‘Accumulibacter phosphatis’ (herein Accumulibacter), and is enriched under cyclic feast (anaerobic, carbon-rich) and famine (aerobic, carbon-poor) regimes (Wentzel et al., 1986; Comeau et al., 1986). Intracellular polyphosphate (polyP) reserves are hydrolyzed under anaerobic conditions to provide energy for uptake and storage of short chain fatty acids, and in the subsequent aerobic phase PAOs uptake phosphate for PolyP replenishment (Comeau et al., 1986; Guisasola et al., 2004) In addition to aerobic P uptake, a select subset of PAOs, termed denitrifying PAOs (DPAOs), are also capable linking P uptake to nitrate (NO_3_) or nitrite (NO_2_) reduction under denitrifying (“anoxic” in environmental bioprocess parlance) conditions (Gao et al., 2017). The activity of DPAOs over NO_2_ is particularly interesting when the goal is to integrate EBPR with the emerging energy efficient shortcut nitrogen (N) removal bioprocesses, where NO_2_ (not NO_3_) is the key intermediate (Gao et al., 2014). From a process standpoint, DPAOs offer an intriguing opportunity to couple N and P removal while decreasing carbon and energy requirements. However, we and others have documented unusually high production of the undesirable potent greenhouse gas nitrous oxide (N_2_O) in denitrifying EBPR bioprocesses (Zhou et al., 2012; Wisniewski et al., 2018). Underlying mechanisms for N_2_O emissions in DPAO-enriched bioprocesses, and associated mitigation strategies, are poorly understood. In particular, while Accumulibacter physiology and patterns of gene expression under anaerobic/aerobic conditions have been reasonably well studied (Comeau et al., 1986; Oehmen et al., 2004; Oyserman et al., 2016), research on anoxic (denitrifying) PAO activity, gene expression patterns, and ecological interactions, especially when NO_2_ (instead of NO_3_) is the electron acceptor, is quite limited (Gao et al., 2019).

Denitrification is a modular pathway in which NO_3_ is enzymatically converted to N_2_ via step-wise reduction by nitrate reductase (*napAB* or *narG,* NO_3_ to NO_2_), nitrite reductase (*nirS* or *nirK,* NO_2_ to nitric oxide [NO]), nitric oxide reductase (*norBC* or *norZ,* NO to N_2_O), and nitrous oxide reductase (*nosZ,* N_2_O to N_2_) (Zumft, 1997). These modular denitrification genes can be regulated with a certain degree of independence (Graf et al., 2014), leading in some cases to accumulation of denitrification intermediates such as N_2_O. Recent work has shown that microbes harboring incomplete (truncated) denitrification pathways that lack one or more key denitrification genes are surprisingly prevalent in both engineered bioprocesses and in natural systems (Gao, 2019; Philippot et al. 2011; Anantharaman et al. 2016). Indeed, our previous work on a denitrifying EBPR process highlighted the possibility that flanking non-PAO denitrifying bacteria lacking nitrous oxide reductase genes may contribute to N_2_O generation (Philippot et al., 2011; Anantharaman et al., 2016; Gao et al., 2019). Although these genomic analyses revealed the genetic potential for incomplete denitrification, we concurrently documented a complete denitrification pathway in the dominant PAO population and a prevalent subset of denitrifiers with nitrous oxide reductase in this system. A better understanding of the role of differential patterns of gene expression and potential for segregation of denitrification metabolism in DPAO-enriched bioprocesses is therefore needed to illuminate mechanisms underlying N_2_O accumulation.

In addition to PAOs, Glycogen Accumulating Organisms (GAOs) are commonly detected in EBPR bioprocesses, as they are also enriched under the anaerobic carbon-rich feast/ aerobic carbon-depleted famine regime (McIlroy et al., 2014). Gammaproteobacterial GAOs affiliated with the *Candidatus* ‘Competibacter’ (herein Competibacter) lineage are commonly observed in these processes (Gregory et al., 2002; Oehmen et al., 2007; McIlroy et al., 2014). While GAOs do not accumulate P, they compete with PAOs for fatty acids during the anaerobic feast period. GAOs are thus widely viewed as undesirable competitors to PAOs that negatively affect the P removal efficiency (Stokholm-Bjerregaard et al., 2017). Current interpretations on the potential interactions between the GAOs and PAOs have focused largely on this competition (Stokholm-Bjerregaard et al., 2017). However, recent work indicates that GAOs may not necessarily be problematic to the enrichment of PAOs (Nielsen et al., 2019). Additional work has suggested the potential collaboration between GAOs and PAOs in denitrification via cross-feeding of NO_2_, as DPAOs are not always able to reduce NO_3_ (Rubio-Rincon et al., 2017). Ecological interactions and potential for similar cross-feeding to influence NO and N_2_O fate is currently unknown.

Biodiversity has been reported in both Accumulibacter PAO populations and in *Competibacter* GAO populations. Based on the phylogeny of the *ppk1* gene, PAOs affiliated with the Accumulibacter genus are classified into two types (I and II), and 14 clades (IA to IE and IIA to II-I) (Camejo et al., 2016; Zhang et al., 2016). Similarly, the intragroup identity of the 16S rRNA gene annotated at > 89% suggested the microdiversity of Competibacter GAO populations (McIlroy et al., 2014). While the composition of PAO and GAO populations are known to differ in various EBPR systems and process variations (Albertsen et al., 2016; Rubio-Rincon et al., 2017), it is not well understood how or if clade-level phylogenetic differentiation aligns with emergent phenotypic variation (e.g. capacity for denitrification or propensity for cross-feeding or N_2_O production).

Another long-standing enigma in understanding the microbiology of EBPR bioprocess is: are the flanking (e.g. non-PAO) populations necessary and how is the biodiversity maintained in the system? Although *Accumulibacter* PAO populations have been reported being enriched to an abundance as high as ∼90% (FISH-based quantification) in lab-scale EBPR reactors (Lu et al., 2006), no lab grown isolates are available to support the phenotypic characterization of the Accumulibacter strains (Zhang et al., 2019). A better understanding on the functional niches of the flanking populations in EBPR processes may shed light on the potential metabolic network between the PAOs and the flanking populations and may also provide hints to guide the enrichment or cultivation of the *Accumulibacter* PAO populations (Handelsman et al., 2004; Lawson et al. 2017; Oyserman et al. 2016; He et al. 2015).

The objective of this study was to investigate how DPAOs, GAOs, and flanking microbial populations with complete or truncated denitrification pathways interact to convert NO_2_ to N_2_O in denitrifying EBPR processes. To this end, we used genome-centric metagenome and metatranscriptomic sequencing analyses to characterize highly active populations, patterns of gene expression, and potential metabolic interactions among active populations in a DPAO enriched bioprocess previously shown to generate high levels of N_2_O via incomplete denitrification (Gao et al., 2017, Gao et al., 2019). Our results reveal a novel clade of *Accumulibacter* putatively adapted to high rate denitrifying P uptake, provide evidence that differential PAO and GAO denitrification gene expression in addition to prevalent truncated denitrification pathways in the flanking community underlie N_2_O emissions, and suggest a previously unobserved cooperation between Accumulibacter and Competibacter via cross-feeding of the denitrification intermediate NO.

## Results

### Denitrifying EBPR Process Characterization

We operated a lab-scale (12 L working volume) denitrifying EBPR sequencing batch reactor fed with synthetic municipal wastewater with cyclic anaerobic/ anoxic phases and a short aerobic polishing phase for N and P removal over NO_2_ for over 3 years. Operational and performance characteristics were described previously (Gao et al., 2017). Briefly, during each cycle, acetate or propionate was dosed to initiate the anaerobic phase, and NO_2_ was dosed at the start of the anoxic phase to simulate effluent from an upstream nitritation reactor. We documented high rate and stable N and P removal accompanied by a substantial production of N_2_O via incomplete denitrification (60-80% of the influent N). During steady-state operation, we chose representative cycles with different primary carbon sources (acetate or propionate) as feed to profile gene expression across different redox conditions. A summary of the conversion of the key C, N and P constituents in this system during these cycles is provided in Figure 1 and Table 1. Regardless of the carbon source, COD was rapidly consumed. COD consumption rates were not significantly different for acetate compared to propionate (ANOVA p-value>0.05). However, we observed higher N removal rate (ANOVA p-value=0.004) and higher N_2_O production (ANOVA p-value=0.033) when propionate was fed as the primary carbon source. The P uptake rates over NO_2_ were comparable (ANOVA p-value>0.05) between the two carbon sources; however, when propionate was supplied as the external carbon source (electron donor), the P-release/C-uptake value (ANOVA p-value=0.016) and overall P-removal efficiency (ANOVA p-value=0.021) were significantly higher. To profile gene expression patterns during SBR operation, we selected a representative cycle for each carbon source to monitor key denitrification genes *nirS* and *nosZ* via RT-qPCR (Table S2). Based on the RT-qPCR results, a single sample in each redox condition (six in total) was selected for metatranscriptomic analyses.

**Figure 1.**
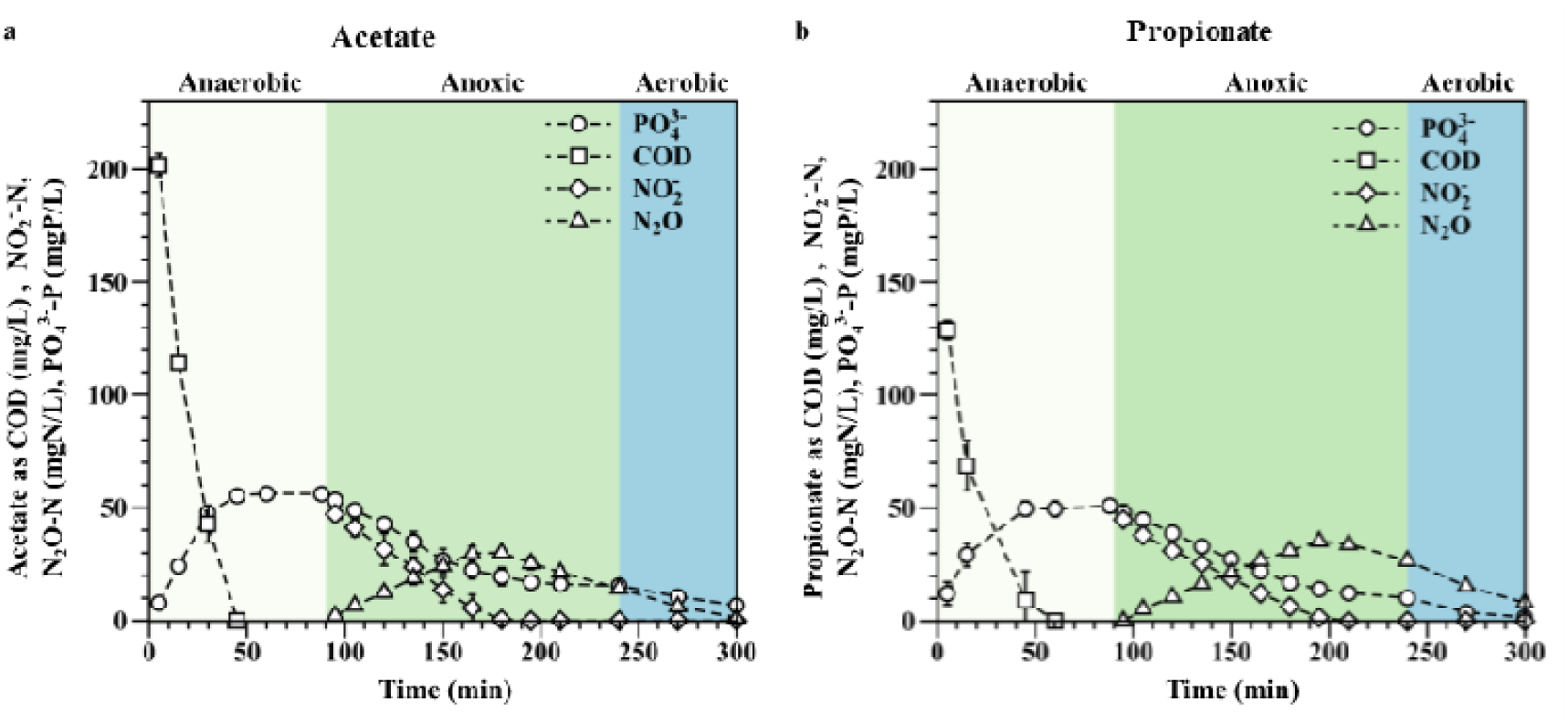
Within-cycle profiles of key C, N and P constituents (acetate/propionate as COD, NO_2_^-^, dissolved N_2_O and PO_4_^3-^) with (a) acetate and (b) propionate feed during RNA sampling. Error bars are based on duplicate measurements.

**Table 1.** N and P removal rates and efficiencies during anoxic and aerobic phases in two typical SBR cycles with different external electron donors (acetate and propionate). Averages and standard deviations are across two cycles for each external electron donor.

|  | Specific anoxic P uptake rate<br>(PO<sub>4</sub><sup>3-</sup> as P)<br>(mg/L•h•MLVV S)<sup>a</sup> | Specific aerobic P uptake rate<br>(PO<sub>4</sub><sup>3-</sup> as P)<br>(mg/L•h•MLVSS)<sup>a</sup> | Specific NO<sub>2</sub><sup>-</sup> reduction rate<br>(NO<sub>2</sub><sup>-</sup> as N)<br>(mg/L•h•MLVSS)<sup>a</sup> | Specific COD removal rate<br>(COD<br>mg/L•h•MLVSS) | P removal (%) | P-release/<br>C-uptake during anaerobic phase<br>(mole/mole) | N<sub>2</sub>O production (%)<sup>b</sup> |
| --- | --- | --- | --- | --- | --- | --- | --- |
| Acetate | 6.6 ± 2.0 | 1.8 ± 0.1 | 6.8 ± 0.2 | 71.8 ± 14.0 | 13.3 ± 12.9 | 0.2 ± 0.0 | 67.7 ± 2.8 |
| Propionate | 6.7 ± 0.5 | 2.6 ± 0.1 | 7.5 ± 0.1 | 68.4 ± 3.4 | 88.2 ± 8.7 | 0.3 ± 0.0 | 80.0 ± 1.5 |

a. NO<sub>2</sub><sup>-</sup> reduction and anoxic/aerobic PO<sub>4</sub><sup>3-</sup> removal rates were calculated based on the linear regression of chemical profiles.
b. Values refer to percent of removed NO<sub>2</sub><sup>-</sup>-N, based on dissolved N<sub>2</sub>O.

**Table 2.**
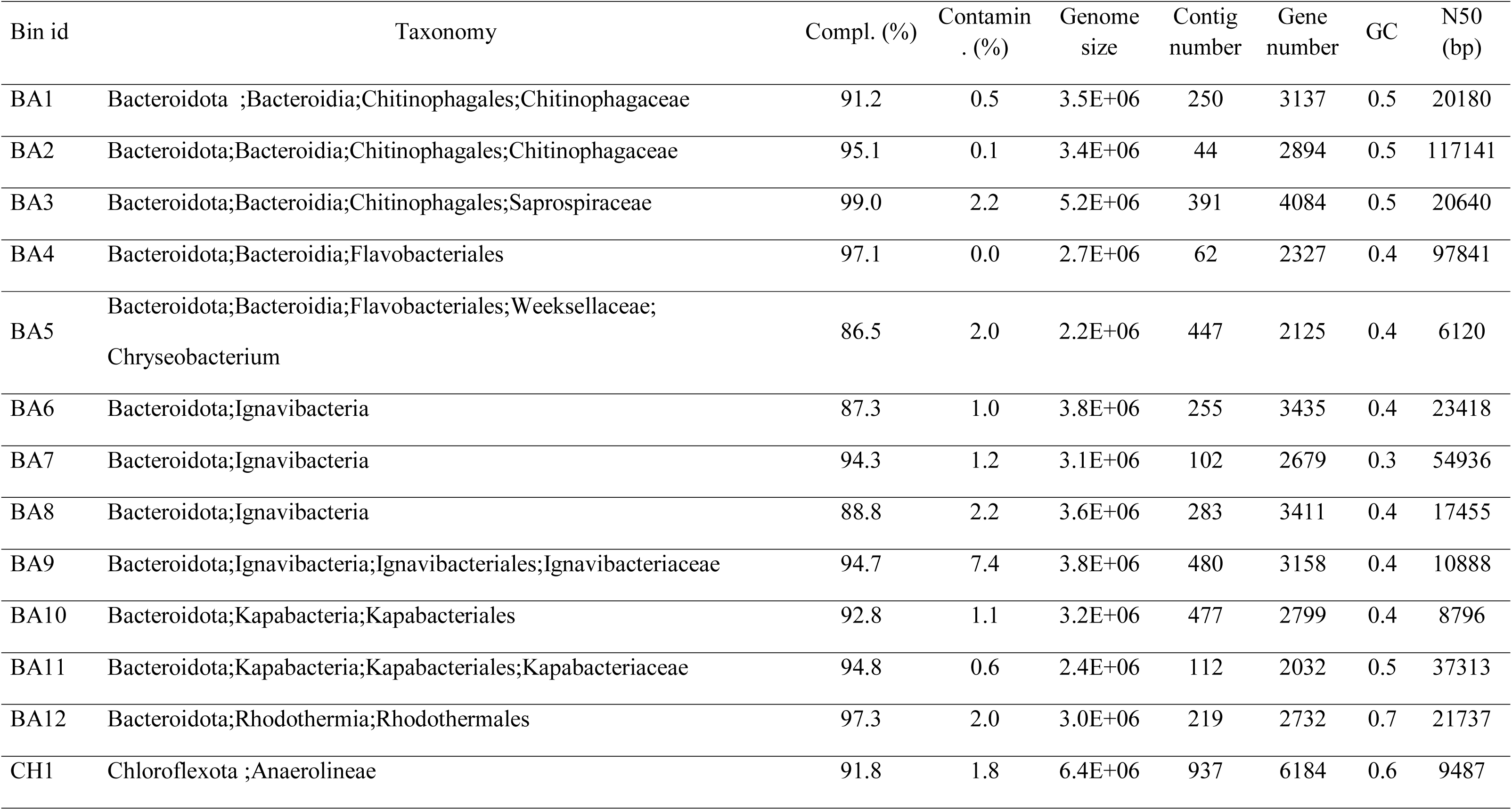
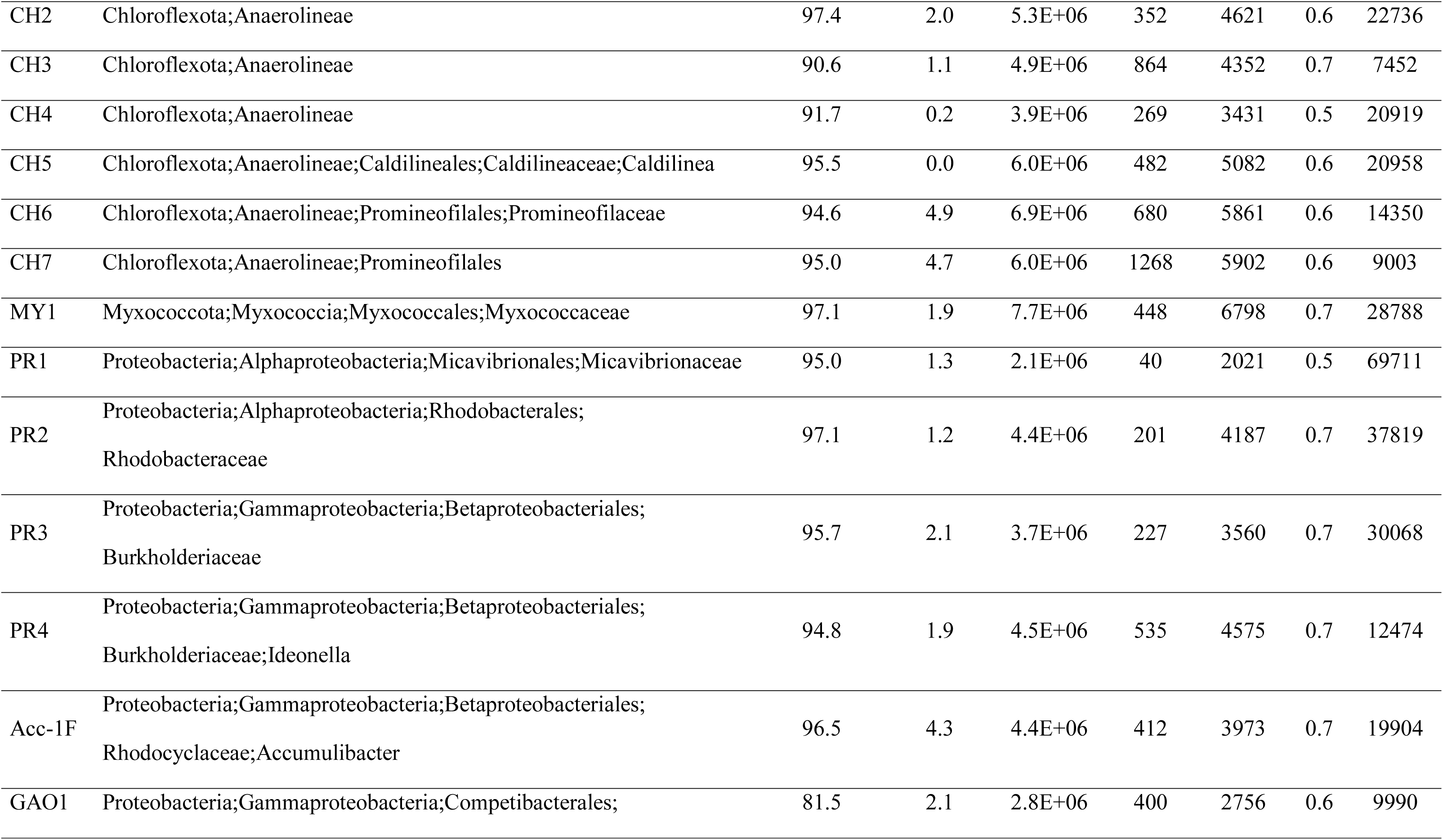
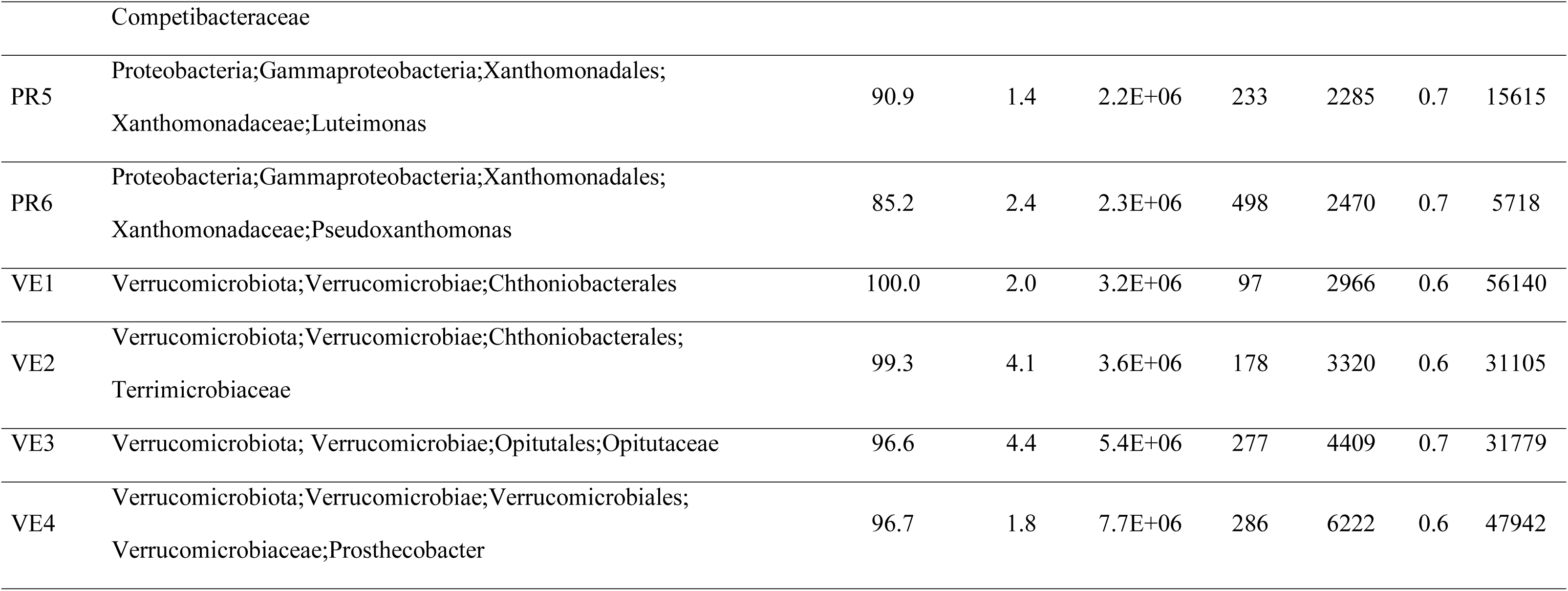
Characteristics of all MAGs obtained in this study, including one novel Accumulibacter genome (Acc-1F), a novel *Competibacter* genome (GAO1), and 30 non-PAO or GAO flanking genomes. The taxonomy affiliation of each MAG is in accord with the Genome Taxonomy Database (GTDB).

It is noteworthy that the P-uptake rate over NO_2_ in this DPAO-enriched consortia was ∼3 times higher than the aerobic P-uptake rate after 3 years of operation (Table 1). In contrast, the anoxic P-uptake /aerobic P-uptake ratio was ∼1.0 in our previous characterization conducted after 7 months of the reactor operation (Gao et al., 2017). NO_3_ and NO_2_ are typically regarded as less efficient electron acceptors for P uptake by PAOs than oxygen, and the denitrifying P uptake rate is generally lower than the aerobic P uptake rate (Kern-Jespersen et al., 1993). The highest ratio between the anoxic P-uptake rate and the aerobic P-uptake rate reported in literature is 0.8:1 (Hu et al., 2002). The elevated activity of P-uptake over NO_2_ coupled to high N_2_O generation as observed in this study is therefore particularly interesting and suggests a strong adaption in this DPAO-enriched biomass to denitrifying conditions. This led us to hypothesize the selection for novel PAO genotypes in this system that correspond with this unique phenotype of proficient NO_2_ utilization.

### PAO microdiversity, identification of a novel Accumulibacter clade, and potential for clade segregation between aggregate size fractions

To profile strain level Accumulibacter diversity, we employed Accumulibacter-specific *ppk1* cloning and sequencing (35 gene sequences), coupled to *in silico* extraction of *ppk1* genes from assembled metagenomic contigs (13 unique gene sequences) and from an Accumulibacter metagenome assembled genome (MAG) described below (Acc-IF). Phylogenetic analysis of the *ppk1* genes (Figure 2) in this DPAO consortia indicated that the Accumulibacter populations were all of Type I, and affiliated with five clades: IA, IB, IC, ID and a distinct new clade, herein termed clade IF. The identity between the *ppk1* genes in this distinct new clade and the *ppk1* genes in the other clades is less than 89.0%. Among the 952 *ppk1* genes downloaded from NCBI, only one *ppk1* gene (KF772928.1) robustly clustered with the clade IF *ppk1* genes obtained in this study, and the highest sequence identity was ∼99.9%. No publicly available genomes affiliate with this novel clade, and to our knowledge, no enrichment cultures nor phenotypic or genotypic characterizations have been described in the literature.

**Figure 2.**
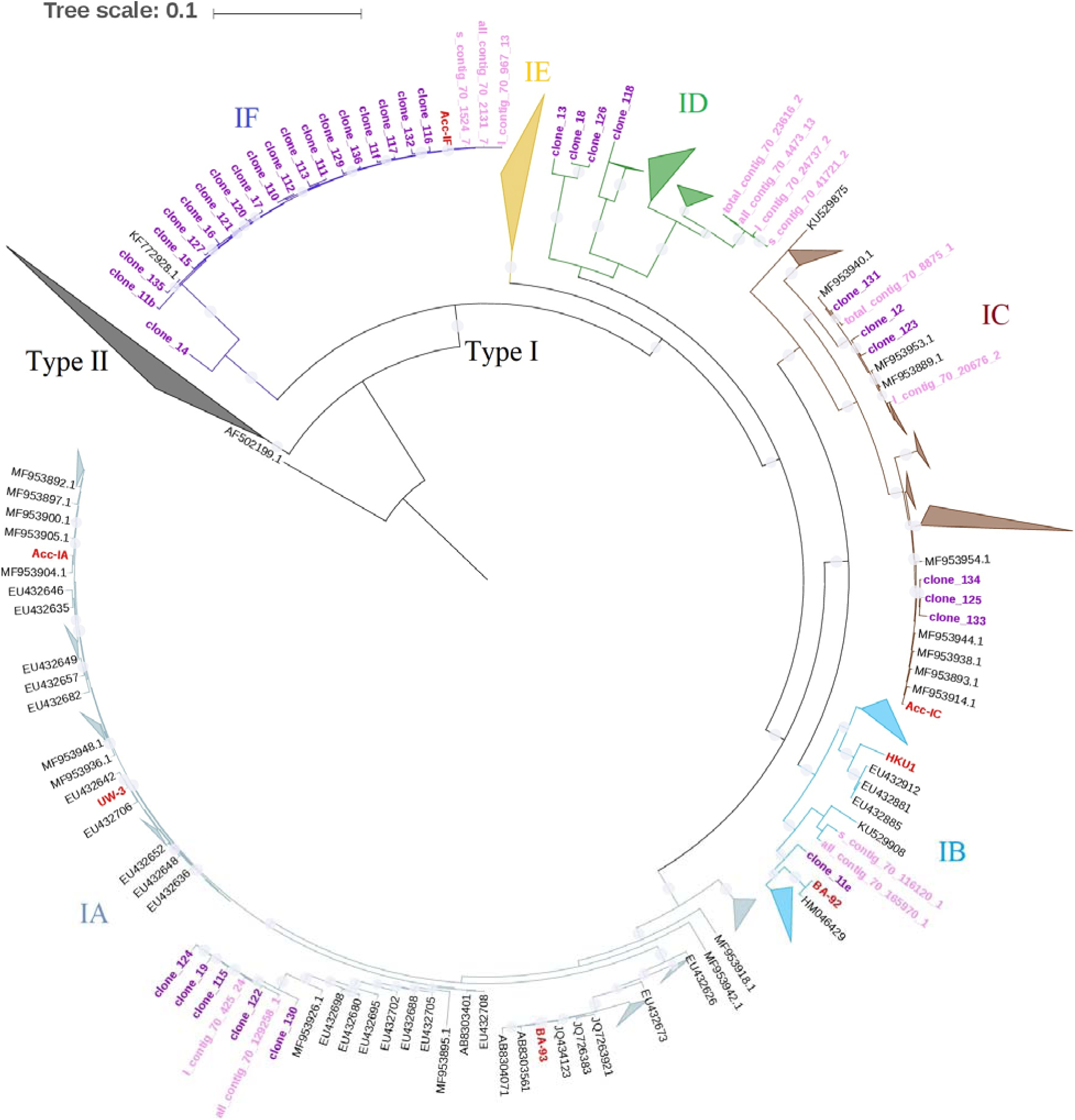
Maximum likelihood phylogenetic tree of Accumulibacter *ppk1* gene sequences (1007 bp fragment) identified in this denitrifying EBPR consortia. Clades IA, IB, IC, ID, and IE are labeled in grey, blue, brown, green and yellow, respectively; and the proposed clade IF is labeled in dark blue. The tree includes *ppk1* genes extracted from 4 available Type 1 genomes and 3 draft genomes from this reactor (sequence names in red; ACC-IF obtained in the study, ACC-IC and ACC-IA from our previous work **(Gao et al., 2019)**); *ppk1* gene sequences obtained via cloning and sequencing (in magenta); *ppk1* gene sequences identified in metagenomic contigs (in pink); and *ppk1* reference gene sequences from NCBI (in black). The *Horocycles tenuis ppk1* gene (AF502199.1) was applied as the outgroup. Type II *ppk1* gene cluster is collapsed in this figure as no *ppk1* gene identified in this DPAO consortia was of Type II, and some sub-clusters of the type I *ppk1* genes were also collapsed to support a clearer visualization of the phylogeny. NCBI accession numbers for Accumulibacter draft genomes BA-93, BA-92, HKU1 are GCA_000585075.1, GCA_000585055.1, GCA_000987395.1. The UW-3 genome was downloaded from JGI Integrated Microbial Genomes (IMG genome ID 2687453699). The accession number of the two Accumulibacter draft genomes Acc-IA and Acc-IC previously reconstructed from the same reactor were PHDR00000000 and PDHS00000000.

While we operated this system for suspended growth (floccular) biomass, granules were formed in this denitrifying EBPR reactor without intentional selection. To understand the potential for population segregation between aggregate types (large granules versus smaller flocs), relative abundances of Accumulibacter clades in granules and in the suspended biomass were evaluated according to the DNA-RPKM values calculated from metagenomic sequencing analyses (Figure S1). The DNA-RPKM values were calculated by mapping the metagenomic DNA reads of the larger sludge particulates (diameter ≥ 350 µm) and the metagenomic DNA reads of the smaller sludge particulates (diameter < 350 µm) to the *ppk1* genes of the five clades, respectively. We found that Accumulibacter clade IA was significantly enriched in larger sludge particulates (or in granules), while the Accumulibacter *ppk1* genes of clade IF and clade IC were of higher ratio in smaller sludge particulates (suspended biomass). Accumulibacte*r ppk1* genes in clades IB and ID were of relatively low abundance in the community.

### Genome-centric profiling of the microbial populations in the DPAO consortia

We assembled 32 high quality (completeness >85%, contamination <5%) MAGs from this denitrifying EBPR process via analysis of shotgun metagenome data (summarized in Table 2). These 32 MAGs phylogenetically affiliate with the phyla Proteobacteria (8), Myxococcota (1), Chloroflexota (7), Verrucomicrobiota(4) and Bacteroidota (12) (taxonomic affiliations are in accord with the Genome Taxonomy Database (Parks et al., 2018). MAG phylogeny is shown in Figure 3. 21 out of these 32 MAGs were significantly divergent from currently available genomes, being annotated with an ANI <85% to currently available reference genomes (see Table S1 for accession numbers for publicly available reference genomes that are of the highest ANI value to each of the 32 MAGs). We recovered a single high quality Accumulibacter MAG (Acc-IF, affiliated with the proposed clade IF) and Competibacter MAG (GAO1) from this analysis. Both of these MAGs represent genotypes distinct from the currently available Accumulibacter and Competibacter genomes. The highest ANI between Acc-IF and 24 publicly available Accumulibacter reference genomes is 82% (Figure S2), and the highest ANI between GAO1 and 19 Competibacter reference genomes is 78% (Figure S3).

**Figure 3.**
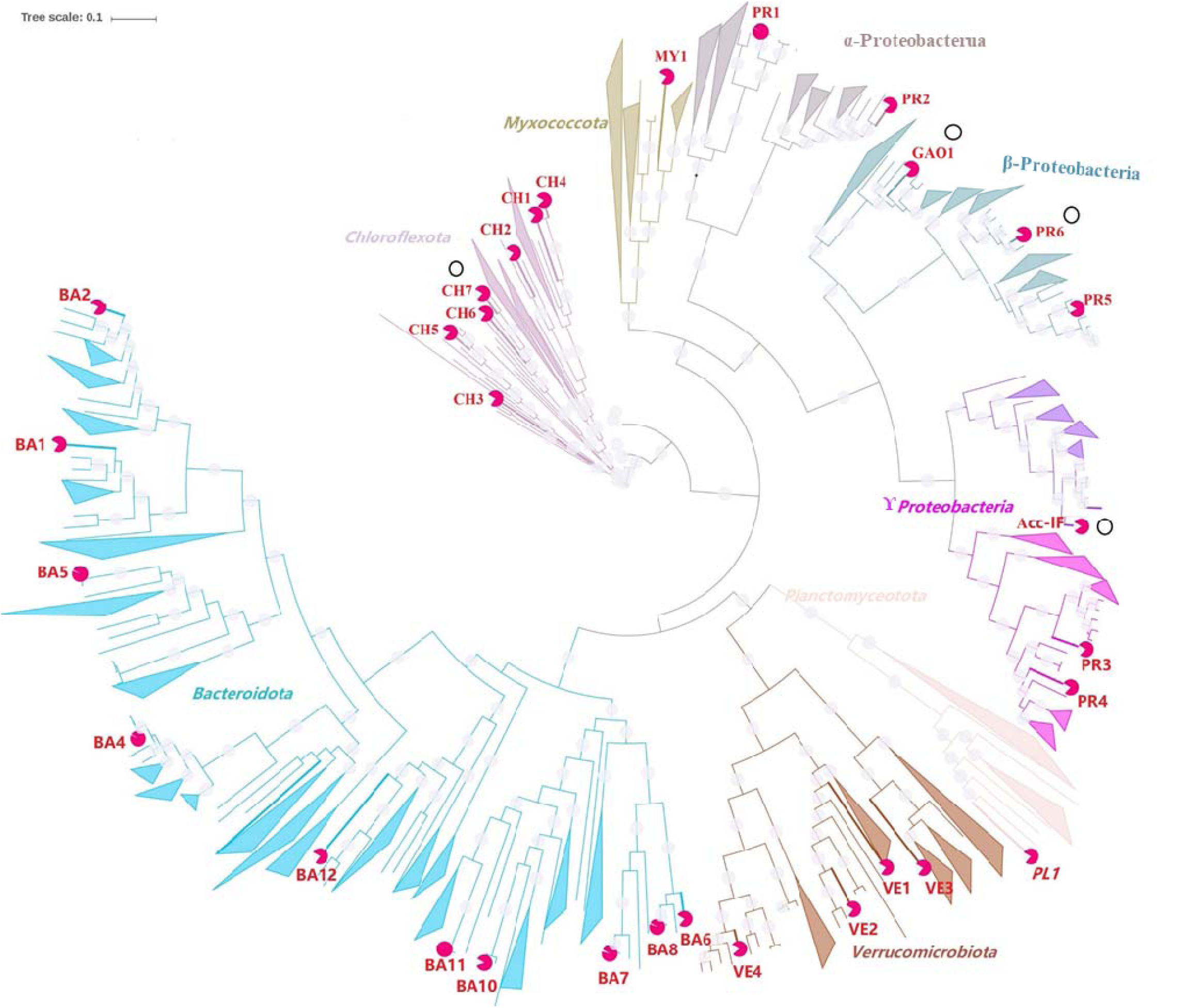
Maximum likelihood tree of the 32 MAGs and phylogenetically closely related reference genomes. Red circles at the end of branches represents the highest ANI between each MAG and the closest publicly available reference genome. The four MAGs of the highly abundant and active populations Acc-IF, GAO1, CH7 and PR6 are highlighted in this figure with a black circle positioned near their label names. Branches of the phylogenetic tree are colored according to the taxonomy affiliation. The tree was constructed using RAxML with 100 bootstraps based on a set of 120 concatenated universal single-copy proteins (Parks et al., 2018).

We estimated relative abundances of the microbial populations corresponding to each of the 32 MAGs based on the DNA-RPKM values calculated from DNA reads mapping, and the associated transcriptional activity (relative gene expression) was estimated based on the RNA-RPKM values calculated from mRNA reads mapping. Results are shown in Figure 4. The RNA-RPKM values of each MAG were averaged across the data at the 6 time points, and the DNA-RPKM values of each MAG were averaged across the values calculated in larger granules, smaller flocs and in the total biomass. As has been summarized above, phylogeny of the *ppk1* genes suggested the presence of Accumulibacter populations in five distinct type I clades (IA, IB, IC, ID and IF). However, only one Accumulibacter MAG with ∼96% completeness and ∼4% contamination (Acc-IF, affiliated with the proposed clade IF) was recovered through the draft genome binning efforts in this study. We previously recovered composite Accumulibacter MAGs from this reactor affiliated with clades IA and IC (Gao et al., 2019). Populations of these two clades were also still present in this reactor, based on *ppk1* gene cloning/ sequencing and analysis of metagenomic contigs (Fig. 2), but were not sufficiently abundant to re-assemble MAGs. To facilitate the investigation of the abundance and the transcriptional activities of the diverse Accumulibacter populations in this reactor, together with the one Acc-IF MAG recovered in this study, we therefore included these two reference Accumulibacter MAGs in clade IA and clade IC as well as a reference clade IB Accumulibacter MAG of clade IB downloaded from NCBI (Skennerton et al., 2014) in the genome-centric transcriptional analyses. No reference genome of Accumulibacter clade ID is available, so this clade was not included in our downstream analysis. Taken together, contigs from the 32 MAGs and the three reference *Accumulibacter* MAGs accounted for 45 ± 2.6% of the total DNA reads and 70.8 ±3.8% of the total mRNA reads.

**Figure 4.**
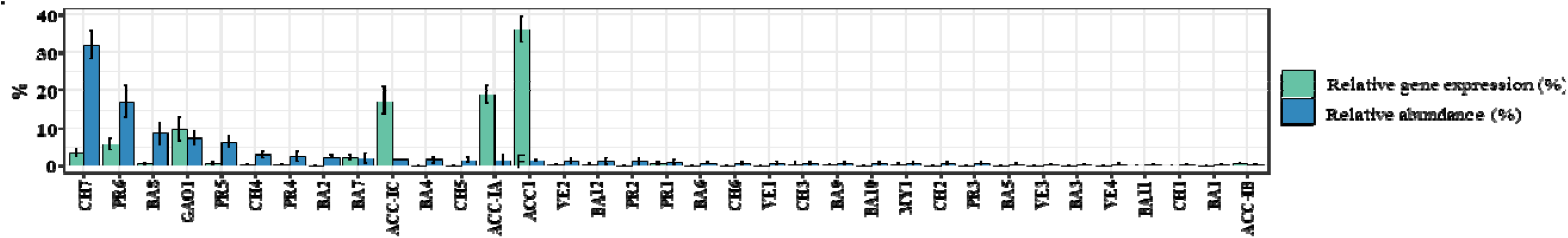
Relative abundance and relative gene expression of 32 MAGs recovered from the denitrifying EBPR reactor in this study and three Accumulibacter draft genomes (ACC-IA [Genbank accession number: PHDR00000000], ACC-IC [Genbank accession number: PDHS00000000] and ACC-IB [Genbank accession number: GCA_000585055.1]). ACC-IC and IA MAGS (affiliated with Accumulibacter clades IC and IA) were recovered previously from this reactor (Gao et al., 2019). For comparison, ACC-1B (affiliated with Accumulibacter clade IB) was included due to identification of *ppk1* genes affiliated with clade IB from cloning and sequencing and from metagenome contig analysis. Draft genomes are ordered based on their genome abundance. Abundance and gene expression estimates are based on DNA-RPKM and RNA-RPKM values of DNA and mRNA reads mapped to each MAG, respectively. Values are averaged across all metagenomic and metatrancriptomic sequencing analyses, with standard deviation indicated by error bars.

Interestingly, as shown in Figure 4, the populations detected with the highest relative abundance (CH7, PR6) were not the ones annotated with the highest transcriptional activities (Acc-IF, Acc-IA and Acc-IC). The metatranscriptome data indicated that Acc-IF was the most transcriptionally active population in the community. Acc-IF alone accounted for 41% of the mRNA reads mapped to the 32 MAGs and the three reference Accumulibacter genomes, while its abundance based on the DNA reads mapping was only ∼2%. The other two highly active Accumulibacter populations represented by Acc-IC and Acc-IA, respectively, accounted for 22% and 16% of the mRNA reads mapped, respectively, but were also present at the timepoint of sampling at relatively low abundance (∼1%). On the other hand, the two flanking populations PR6 and CH7, to which only 3-5% of the mRNA reads were mapped, represented 15% and 35% of the metagenome DNA reads, respectively. The stark variation between abundance (based on metagenomic sequencing) and transcriptional activity (based on metatranscriptomic sequencing) is surprising. Because they reflect gene expression patterns rather than genomic potential, RNA-seq based transcriptional analyses are commonly thought to more accurately reflect functional activity than the DNA-seq based metagenomic analyses (Oyserman et al., 2016). The top 6 transcriptionally most active populations (Accumulibacter Acc-IF, Acc-IC, Acc-IA, Competibacter GAO1, CH7 and *Pseudoxanthomonas* PR6) accounted for 92% of the mRNA reads mapped to the 32 MAGs and the three reference Accumulibacter genomes. By comparison, the transcriptional activities of the populations represented by the other 28 MAGs and the Accumulibacter reference genome Acc-IB were very weak.

### Truncated denitrification pathways and differential gene expression as controls on N_2_O production

Substantial accumulation of N_2_O (60-80% of N fed to the system) from NO_2_ reduction was detected in this denitrifying EBPR bioprocess. To identify putative NO_2_ reducers and N_2_O producers in the community and to understand the underlying mechanisms for N_2_O accumulation, we queried all MAGs for the presence/absence and the expression of the core denitrification genes *napAB*, *narG*, *nirS*, *nirK*, *norBC*, *norZ*, and *nosZ*. As summarized in Figure 5, among the 32 MAGs recovered in this study, 2 MAGs (BA7 and VE3) contained no denitrification genes, and only 3 MAGs (Acc-IF, BA4 and MY1) harbored a complete denitrification pathway (that is, genomic machinery for reduction of NO_3_ to N_2_). In contrast, a large proportion (27 out of 32) of recovered MAGs harbored truncated (incomplete) denitrification pathways that lacked one or more key denitrification genes. This group included the single putative GAO (Competibacter, GAO1) that harbored genes for nitrate, nitrite, and nitric oxide reductase (*narG*, *nirS*, and *norB*), but lacked nitrous oxide reductase (*nosZ*). Taken together, these results demonstrate a high prevalence of genomes that harbor incomplete denitrification pathways---that is, that harbor genomic capacity to catalyze at least one step of the reduction of NO_3_^-^ to N_2_, but not genes encoding the complete denitrification pathway. This result is consistent with our metagenomic analyses in this system at an earlier stage of operation that also demonstrated highly prevalent incomplete denitrifiers (Gao et al., 2019). However, these results demonstrate genomic potential, but not expression, of denitrification genes. To assess how genomic potential relates to gene expression and associated reactor phenotype in this system, we employed metatranscriptomic sequence data to investigate the expression of genes related to N metabolisms.

**Figure 5.**
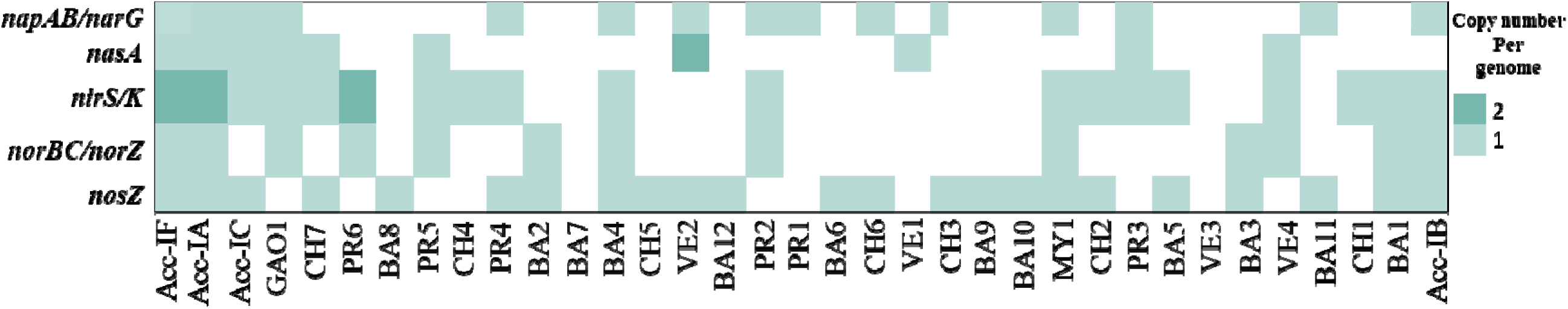
Presence (Warnecke et al.) or absence (white) of core denitrification genes in 32 MAGs and the three reference Accumulibacter genomes. Acc-IF (recovered in this study), Acc-IA, Acc-IC (recovered previously from this system) and Acc-IB are Accumulibacter-affiliated MAGs; GAO1 is the one *Competibacter* affiliated MAG. *napAB*: periplasmic nitrate reductase; narG: nitrate reductase; *nasA*: nitrate transporter; *nirS/K*: nitrite reductase; *norBC*: respiratory nitric oxide reductase; *norZ*: nitric oxide reductase; *nosZ*: nitrous oxide reductase.

To explore whether differential gene expression may be linked to the unusually high levels of N_2_O production we observed in this reactor, we first compared expression of *nirS* and *nosZ* in both the overall community and by Accumulibacter populations. Denitrification gene expression based on metatranscriptomic sequencing analyses is summarized in Figure 6. The *nirS* gene (RNA-RPKM value of 3351), not the *nirK* gene (RNA-RPKM value of 24), was the dominant nitrite reductase gene in this consortium. Consistent with the overall transcriptional activity, ∼93% of the denitrification gene transcripts also mapped to the top 6 transcriptionally active MAGs (Acc-IF, Acc-IA, Acc-IC, GAO1, CH7, and PR6). Our analysis revealed strongly elevated transcriptional activity of *nirS* (nitrite reductase) compared to *nosZ* (nitrous oxide reductase) in both the overall community and in the three highly active Accumulibacter MAGs. ∼90% of the *nirS* gene transcripts were associated with the three Accumulibacter MAGs Acc-IF (∼45%), Acc-IA (35%) and Acc-IC (10%), suggesting that denitrifying PAOs were the major denitrifiers utilizing NO_2_. Similarly, *nosZ* gene transcripts were also predominantly associated with the Accumulibacter MAGs: Acc-IF (∼37%) and Acc-IA (33%). However, the expression levels of the *nirS* gene versus the *nosZ* gene were highly imbalanced in the Accumulibacter populations, and the RNA-RPKM values of the *nirS* gene were ∼7 times larger than that of the *nosZ* gene. Similarly, under all conditions, the global expression of *nirS* (average RPKM=2832.3±788.0) was significantly higher compared to *nosZ* (average RPKM=775.7±199.4) (ANOVA p-value=0.0001). We previously observed that the Acc-IC genome encoded no nitric oxide reductase gene and an incomplete *nosZ* with stop codons within the ORF (Gao et al., 2019). Consistent with this observation, the RPKM value for Acc-IC associated *nosZ* was less than 3 for all the 6 sampling points, suggesting a non-functional *nosZ* gene in the Acc-IC population in our reactor. Taken together, the imbalance in expression levels of *nirS* versus *nosZ* genes in Accumulibacter draft genomes suggests that Accumulibacter is likely an active NO_2_ reducer (as evidenced by high denitrifying P uptake, Table 1), and the low transcription activity of *nosZ* gene in Accumulibacter populations may have induced the N_2_O accumulation.

**Figure 6.**
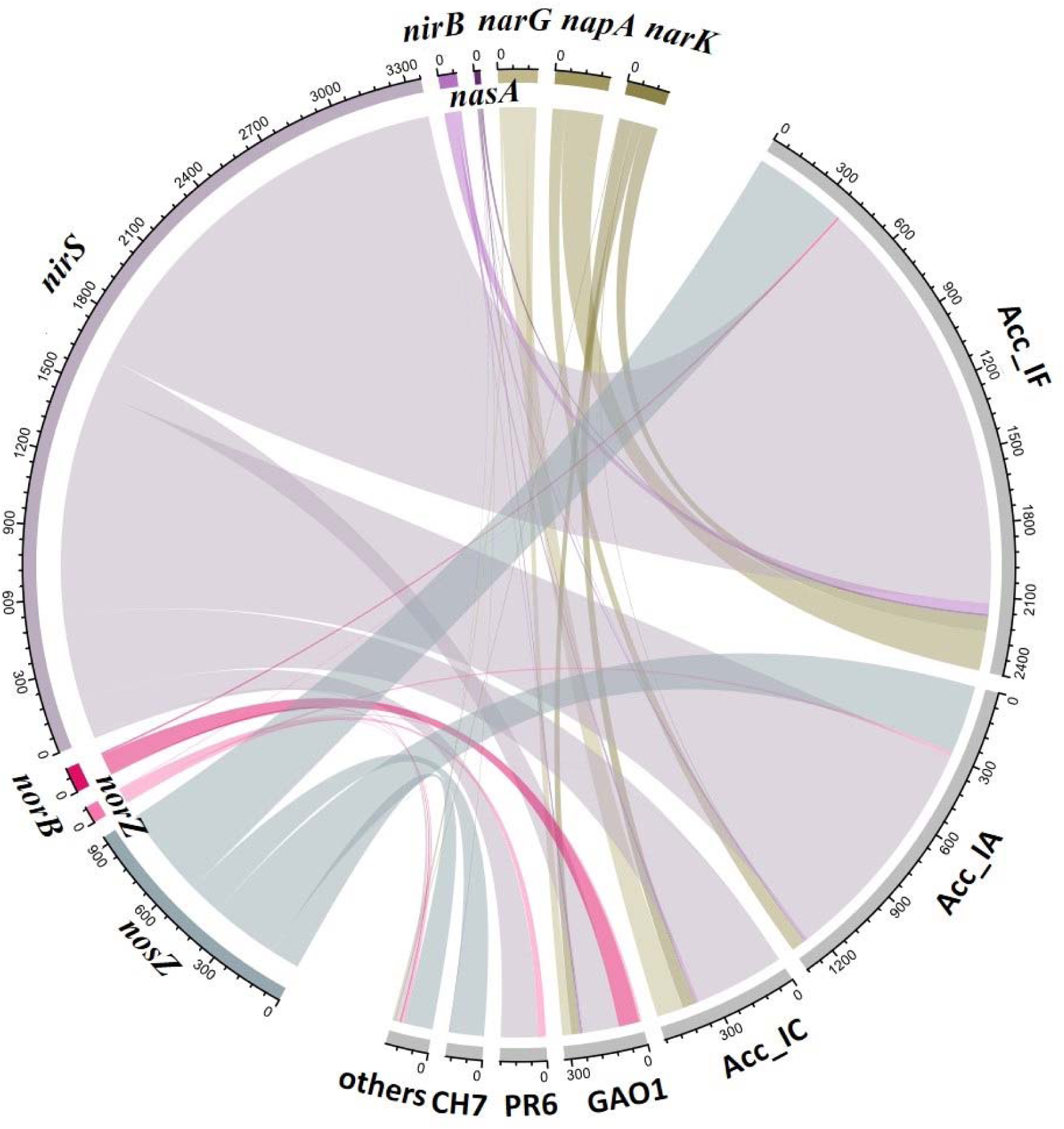
RNA-RPKM values of core denitrification genes encoding in the overall microbial community (left, organized by genes) and in the six MAGs of the highest expression activity (right, organized by genomes) under anoxic conditions. *nirK* expression levels were >100 fold lower than *nirS*, and are therefore not shown. Colors represent different genes, and the width of each ribbon represents the RNA-RPKM value averaged across the RNA-RPKM values of each gene under the anoxic period applying the two carbon sources: acetate and propionate.

### Potential cooperation between GAOs and PAOs and putative N_2_O producers and consumers

Our analysis revealed a surprising potential segregation of denitrification metabolism and cooperative cross-feeding between the PAO (Accumulibacter) populations and the GAO population. While the three Accumulibacter MAGs accounted for the vast majority of nitrate reductase (*narG* and *napAB),* nitrite reductase (*nirS)*, and nitrous oxide (*nosZ*) gene transcripts, ∼93% of the *norB* gene transcripts were associated with the *Competibacter* MAG GAO1 (Figure 6). This suggested that GAO1 was the dominant NO reducer and N_2_O producer in this system. It should be noted that expression of NOR (*norB* or *norZ*) was low compared to both *nirS* and *nosZ.* Although *norB* genes were annotated in both Acc-IA and Acc-IF, the *norB* genes were not highly expressed in these *Accumulibacter* MAGs. Interestingly, similar to several other publicly available Accumulibacter genomes (Flowers et al., 2013; Camejo et al., 2019; Speth et al., 2016), a nitric oxide reductase gene (*norB* or *norZ*) was not identified in Acc-IC. The strong differences in *norB* versus *nirS* gene expression between PAO and GAO populations suggest the possibility for metabolic segregation of denitrification and cooperative cross-feeding of NO between Accumulibacter and Competibacter.

As discussed above, in addition to dominating *norB* transcriptional activity in this community, GAO1 also lacks a *nosZ* gene and is therefore incapable of N_2_O reduction. PR6 is similarly a putative N_2_O producer, based on the presence of an incomplete denitrification pathway lacking genomic potential for N_2_O reduction (Figure 5). It is therefore likely that GAO1 and PR6 are key N_2_O producers in this denitrifying EBPR system, as both GAO1 and PR6 lack genomic potential for N_2_O consumption, while they account for the majority of the transcriptional activity of the *norB* gene and the *norZ* gene, respectively. Conversely, our analyses suggest that the flanking population CH7 may be an important N_2_O consumer in this community, due to the fact that expression of *nosZ* in CH7 was higher than that of *nirS*, particularly under anoxic and aerobic conditions (Figure S4). Interestingly, no nitric oxide reductase genes were annotated in CH7.

### Transcriptional activity of genes involved in acetate and propionate utilization

Based on comparative kinetic analyses between SBR cycles utilizing acetate and propionate as the primary carbon source (see Table 1), differences in terms of P removal efficiency (as well as P-release/C-uptake) and N_2_O production were observed. To understand the impact of and competition for primary carbon sources among bacterial taxa in this denitrifying EBPR system, we explored the genome-specific expression of two genes responsible for the activation of acetate, acetyl-coenzyme A synthase (*acs*, high affinity) and acetate kinase (*ackA*, low affinity), and the gene related to propionate activation, propionyl-CoA synthetase (*prpE*) (Figure 7). Compared with the genomic potential for propionate activation, the capability for utilizing acetate was more widespread among the MAGs. Only 2 out of the 32 MAGs lacked acetate activation genes, while only 6 harbored propionate activation gene (Figure 7a). As summarized in Figure 7b, over 95% of the *acs*, *ackA* and *prpE* gene transcripts mapped to the three PAO MAGs (Acc-IF, Acc-IA, Acc-IC) and the one GAO MAG (GAO1), confirming that their ability to efficiently sequester these VFAs during the anaerobic period likely gave these PAOs and GAOs a selective advantage over other members of the community for subsequent NO_2_ denitrification. The expression of the high affinity *ackA* gene was substantially higher (∼5-fold) than the low affinity *acs* gene in the PAO and GAO populations, with the exception that Acc-IA expressed an equal amount of the *ackA* and *acs* gene transcripts, both at low levels. By comparing the expression activity of the *acs*, *ackA* and *prpE* genes, putative variations in the preferred carbon sources between PAOs and GAOs as well as between the three *Accumulibacter* genomes were revealed. On average, Acc-IF, Acc-IA, Acc-IC and GAO1 accounted for 42.0±2.7%, 19.7±1.9%, 26.0±2.9% and 10.9±1.2% of the total *prpE* gene transcripts, respectively. Comparing with the *prpE* gene, the expression of *acs* gene was more evenly distributed, with Acc-IF, Acc-IA, Acc-IC and GAO1 accounting for 35.1±2.5%, 9.2±2.4%, 32.1±5.0%, and 17.1±1.6% of the *acs* gene transcripts, respectively. These results suggest that GAO1 may utilize a higher portion of the primary carbon source when acetate rather propionate was fed into the reactors.

**Figure 7.**
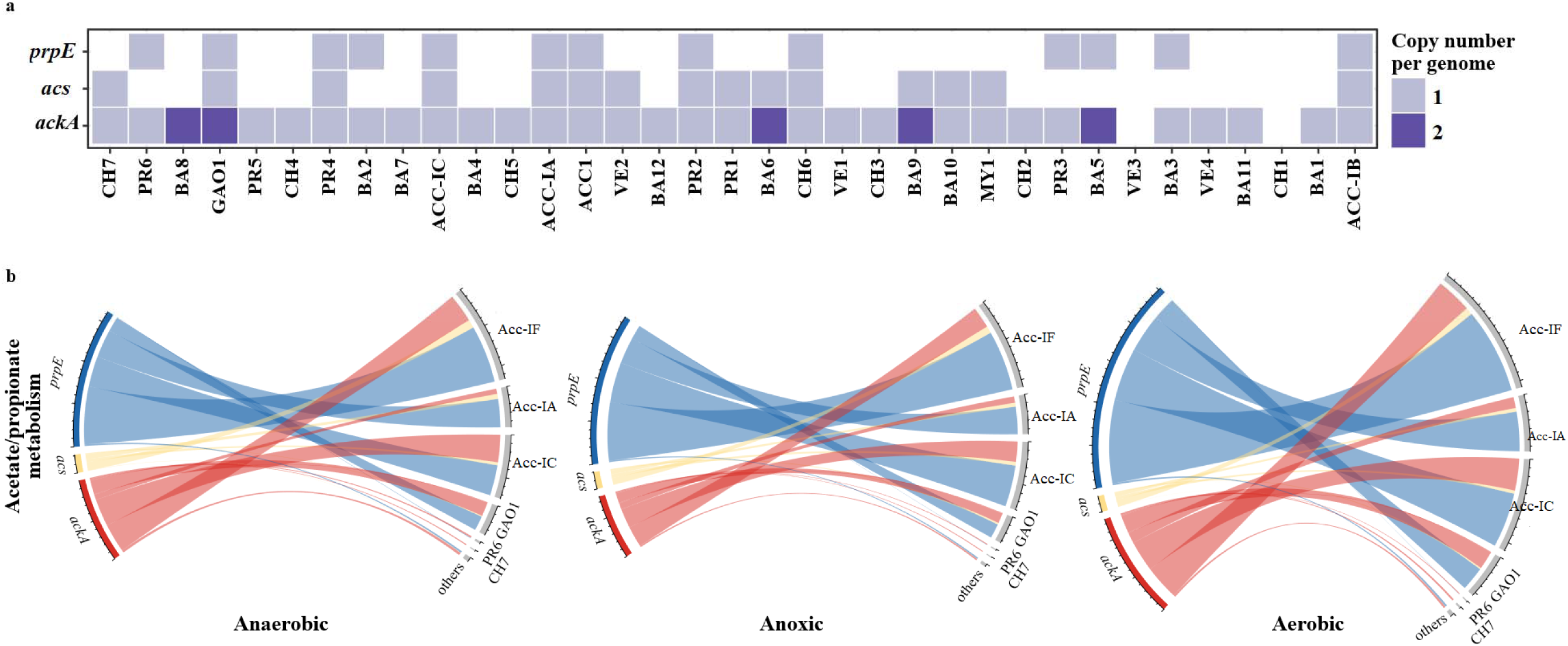
(a) The presence of key acetate/propionate activation genes (acetyl-coenzyme A synthase: *acs*, acetate kinase: *ackA*, and propionyl-CoA synthetase: *prpE*) in the 32 MAGs and three reference *Accumulibacter* draft genomes. Bacterial draft genomes are ordered based on their genome abundance. (b) RPKM values of selected acetate/ propionate activation genes in the overall microbial community (left, organized by genes) and in each MAG (right, organized by genomes) under different redox conditions (anaerobic/anoxic/aerobic). Different colors represent different genes, and the width of the ribbons represent the RNA-RPKM values averaged across the RNA-RPKM values of each gene applying the two carbon sources: acetate and propionate.

### Carbon and Energy Metabolism and Potential Metabolite Exchange among PAOs and Flanking Populations

Figure S6 shows the gene expression profiles and dynamics of major carbon, energy and phosphate metabolic pathways across the top six identified highly active MAGs. Apart from their key functional role in polyP accumulation, as expected, genes involved in both glycolysis and the TCA cycle were annotated as being actively expressed in the three Accumulibacter populations of clade IF, IC and IA, and poly(3-hydroxyalkanoate) polymerase subunits *PhaE/ PhaC* involved in PHA synthesis were also highly expressed in these populations. In addition, all three Accumulibacter populations, especially Acc-IF, were identified as being dominant in the expression of genes involved in EPS synthesis, in the biosynthesis of type IV pilus, in amino acid metabolism, and in the biosynthesis of co-factors like vitamin B12 and Biotin (Figure S5 and Figure S6).

Among the 31 MAGs we recovered representing flanking non-PAO populations, only 3 were identified as being highly transcriptionally active: GAO1 (Competibacter), CH7 (affiliated with the class Anaerolineae) and PR6 (affiliated with the genus *Pseudoxanthomonas)*. These flanking populations were all heterotrophs, with no carbon fixation pathway identified in their genomes. All MAGs expressed genes involved in central carbon metabolic pathways, including the TCA cycle, pyruvate metabolism, pentose phosphate pathway (PPP) and glycolysis/gluconeogenesis (Figure S6). Whereas GAO1 likely utilizes acetate and propionate as its primary carbon source based on gene expression patterns, the expression levels of *acs* and *ackA* genes for acetate activation were comparatively low in both CH7 and PR6. Moreover, CH7 is apparently incapable of utilizing propionate, since no *prpE* gene was identified in its genome. As summarized in Figure S5, 88% of the *vpr* gene transcripts for extracellular protease were mapped to CH7, and 84% gene transcripts for extracellular serine protease associated with PR6. Interestingly, both PR6 and CH7 expressed genes encoding glycoside hydrolases (xylosidase, lysozyme, glucosidase, and isoamylase) and extracellular cellulose binding enzymes that are linked to the breakdown of polysaccharides such as xylan, starch and peptidoglycan (Figure S7).

These results suggest that CH7 and PR6 may scavenge extracellular substances (endogenous organic carbon) excreted by the PAO populations as their primary carbon sources. In addition, the vitamin B12 transporter *btuB* gene and *tonB* gene were highly expressed in PR6 (Figure S5). *TonB* was reported as being essential in importing essential micronutrients across the outer cell membrane (Shultis et al., 2016). PR6 also contributed 33% of the *PilQ* and *PilY1* gene transcripts for type IV pilus biogenesis, and therefore may play a supporting role the three Accumulibacter populations in aggregate (biofilm) formation. Based on patterns of gene expression in carbon, nitrogen, and P cycling, a conceptual schematic of hypothesized metabolic interactions between the Accumulibacter population, GAO1, and the flanking population CH7 in this DPAO-enriched consortium is shown in Figure 8.

**Figure 8.**
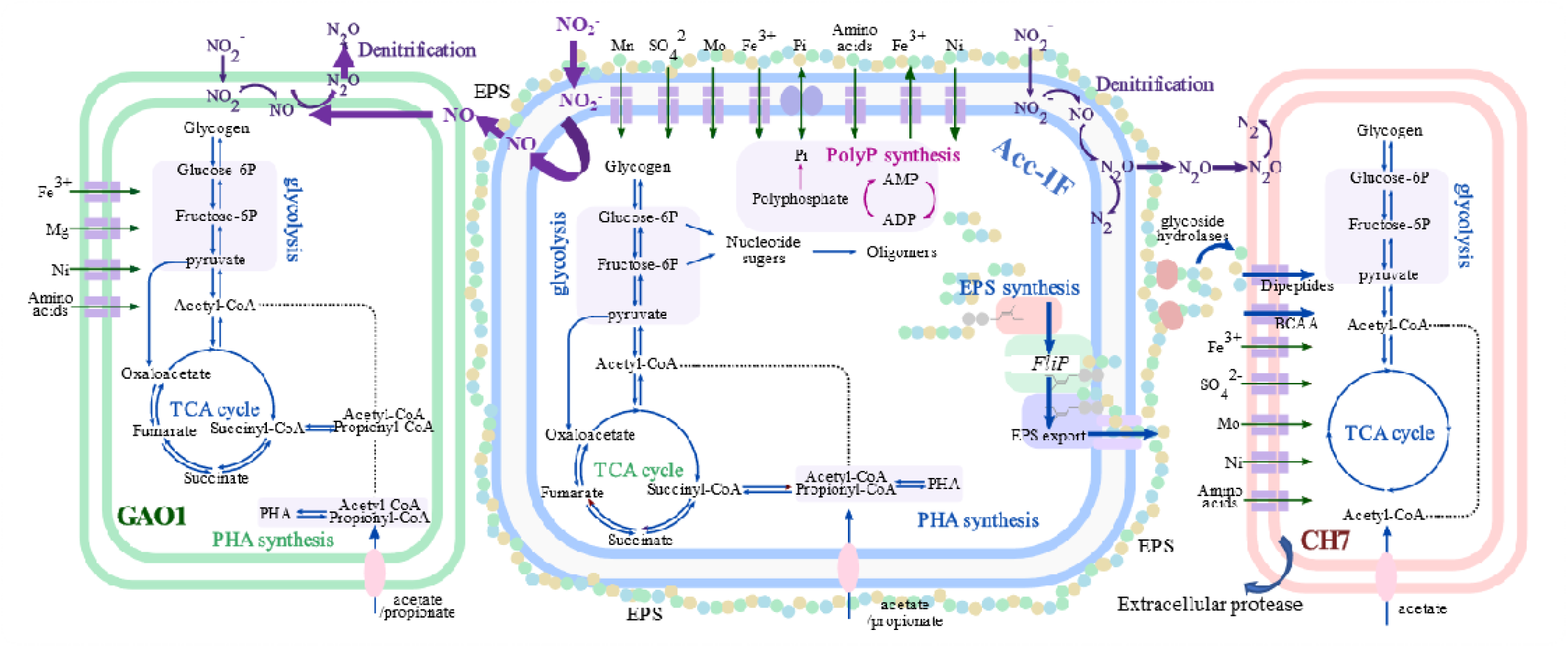
Conceptual schematic of predicted metabolic interactions between Accumulibacter associated PAOs (ACC1, middle), Competibacter associated GAOs (GAO1, left) and CH7 (Haitjema et al.) in the denitrifying EBPR biomass, based on integrated metagenomic and metatranscriptomic sequencing analyses. Carbon metabolic pathways, PolyP synthesis, denitrification is labelled in blue, magenta and purple, respectively. Circles with different colors are used to represent EPS produced by Accumulibacter. Arrows illustrating hypothesized substrate exchange between *Accumulibacter* and CH7 are bolded.

## Discussion

### Segregation of denitrification metabolism between GAOs and PAOs

Denitrification is a modular metabolic process in which NO_3_ or NO_2_ is sequentially reduced to N_2_ via metabolic intermediates NO and N_2_O. While denitrification to N_2_ is commonly assumed to rely on activity of denitrifying taxa with complete denitrification pathways, it is predicted that segregating metabolic processes into different cell types would eliminate inter-enzyme competition and reduce the accumulation of metabolic intermediates. Such segregation has been proposed to be especially beneficial when the metabolic intermediates are toxic and growth-inhibiting (Liljia and Johnson, 2016). In the DPAO-enriched consortia that is the focus of this study, denitrification gene transcripts coupled to genome-resolved metagenomic analyses suggested a potential segregation of NO_2_^-^ and NO reduction into PAO and GAO populations, respectively, as ∼90% of the nitrite reductase gene transcripts were expressed by three co-occurring PAO populations, while ∼93% of the nitric oxide reductase gene transcripts were expressed by the one GAO population.

This hypothesized segregation and cross feeding of NO is in accordance with our previous evaluation on the evolution of the denitrification traits within *Accumulibacter* genotypes: periplasmic nitrate reductase (*napAGH*), cytochrome cd1 nitrite reductase (*nirS*), and the nitrous oxide reductase (*nosZDFL*) were identified as core genes to all type I Accumulibacter populations, while nitric oxide reductase (*norB*) was annotated as a flexible gene for both type I and type II Accumulibacter. Moreover, a nitric oxide reductase gene was not annotated in one *Accumulibacter* MAG Acc-IC recovered from this system (Gao et al., 2019). This finding is surprising given that NO is toxic to most microorganisms at low concentrations, but is consistent with other recent studies that found that only a subset of Accumulibacter genomes harbor complete nitric oxide reductase genes (Flowers et al., 2013; Camejo et al., 2016; Mao et al., 2014). Additional work focused on non-PAO denitrifiers has also documented a puzzling lack of canonical *nor* genes (Grat et al., 2014; Speth et al., 2016; Hallin et al., 2018). Moreover, in an Accumulibacter enrichment culture operated with cyclic anaerobic/ microaerobic conditions, Camejo et al. documented very low expression of *nor* compared to *nirS* and *nosZ* under microaerobic conditions (Camejo et al., 2019). This is consistent with our results, where *nor* expression was much lower than *nirS* and *nosZ* both in the overall community and in Accumulibacter populations. It is also consistent with metaproteomic analyses by Barr et al. (Barr et al., 2016), who detected nitrite and nitrous oxide reductase proteins but no nitric oxide reductase proteins in an Accumulibacter-enriched biofilm, despite presence of a *norZ* gene in the Accumulibacter genome. Camejo et al. speculated that the lack of *nor* in many Accumulibacter genomes may be linked to NO cross feeding and consumption with the flanking non-PAO community (Camejo et al., 2016). Based on genome-resolved metagenomics analyses of a partial nitritation/anammox reactor, Speth et al. speculated that NO might be released into the solution as a denitrification intermediate that might be further removed through aeration or be cross fed to anammox to be further metabolized to N_2_, as few populations annotated with a *nirS/K* gene were also annotated with a *norBC/Z* gene (Speth et al., 2016). Similarly, potential segregation of NO and the NO_2_ reduction among different populations was also suggested based on metatranscriptomic sequencing data of anammox granules (Lawson et al., 2017).

While accumulation of the metabolic intermediates NO_2_ and N_2_O are commonly reported in denitrifying cultures, reports of substantial NO accumulation is rare (Kampschreue et al., 2009; Ge et al., 2012; Hassan et al., 2016; Wertz et al., 2018). Based on the integrated meta-omic analyses here and results from these previous studies, we speculate that the toxicity of NO may promote metabolic segregation of NO reduction into a population that is different from that which reduces NO_2_ to NO. In this DPAO-enriched consortium, patterns of gene expression suggest that PAO populations are the key NO_2_ reducers, while GAOs are the key NO reducers. Such metabolic segregation may be beneficial in eliminating the accumulation of the toxic NO intermediate. This is analogous to the prediction that the segregation of the NO_3_ and NO_2_ reduction reduces the accumulation of NO_2_ (Lilja and Johnson, 2016), which can be inhibitory to many denitrifiers at high concentrations. It should be noted that the potential segregation of NO_2_ and NO reduction in the DPAO-enriched consortia that is our focus is based solely on the gene transcriptional patterns mapped to PAO and GAO MAGs. Further wet-lab experiments are necessary to conclusively demonstrate NO crossfeeding, and more research is needed to identify how common such segregation might be in denitrifying communities. The efforts of Lilja and Johnson (Lilja and Johnson, 2016) with model dual species communities of denitrifiers offers an appearing route to directly test the fitness benefits of metabolic segregation at different steps in the denitrification pathway.

### Mechanisms for N_2_O production by the denitrifying EBPR consortia

Our results suggest a strong role for differential denitrification gene expression in both Accumulibacter populations and in the overall microbial community in controlling N_2_O accumulation in DPAO-enriched consortia. Unlike *norB* gene transcripts (∼93% of which were expressed by the one GAO population), >70% of both the nitrite reductase gene (*nirS*) transcripts and the nitrous oxide reductase gene (n*osZ*) transcripts were associated with the *Accumulibacter* populations. However, the RNA-RPKM values of the *Accumulibacter nirS* genes were ∼7 times higher than that of the Accumulibacter *nosZ* genes. Considering that NOS is responsible for the conversion of N_2_O to N_2_, such differential expression of the *nirS* gene versus the *nosZ* gene in the Accumulibacter populations may limit N_2_O conversion to N_2_ and induce the observed N_2_O accumulation in this and other denitrifying EBPR bioreactors. Such imbalance between *nirS* and *nosZ* gene abundance and expression has also been noted as potential factor leading to N_2_O generation by non-PAO denitrifiers in soils and other environments (Philippot et al., 2009; Schreiber et al., 2012). Another factor that may affect N_2_O accumulation is inter-enzyme competition between the nitrite reductase and the nitrous oxide reductase in the *Accumulibacter* populations. Competition for electrons has been demonstrated among N-reductases, which would cause the accumulation of denitrification intermediates (Pan et al., 2013). If the nitrite reductase has priority over (e.g. outcompetes) nitrous oxide reductase in the utilization of either reducing equivalents or the building blocks for enzyme synthesis, the N_2_O reducing capacity by the host Accumulibacter populations may be limited when NO_2_ is available as the electron acceptor.

### Potential cooperation and competition patterns between PAOs, GAOs, and flanking populations in the denitrifying EBPR biomass

Accumulibacter-affiliated PAOs and Competibacter-affiliated GAOs are two key functional groups in EBPR bioprocesses (McIlroy et al., 2014). We previously recovered a novel *Accumulibacter* MAG in clade IC (Gao et al., 2019) from this denitrifying EBPR consortium. In this study, we recovered a novel Accumulibacter MAG associated with an entirely new clade (proposed as clade IF), as well as novel GAO MAG. The denitrifying EBPR consortia in this study was characterized by the unique feature of significant N_2_O accumulation and highly efficient P-uptake over NO_2_. Recovery of these novel GAO and PAO MAGs is important in expanding our understanding of novel genotype features corresponding to the unusual phenotypes observed in this and other denitrifying EBPR processes (Camejo et al., 2016). In particular, while the proposed Accumulibacter clade IF has not been previously documented in these types of bioprocesses, its exceptionally high transcriptional activity under anoxic conditions and the unusually high P uptake over NO_2_ compared to O_2_ (Table 1) suggest that this clade may be well suited to denitrifying biological P removal. Importantly, differential expression of *nirS* versus *nosZ* also suggest that this clade may be linked to excess N_2_O emissions.

We documented a significantly higher transcriptional activity for Accumulibacter-associated *prpE* over Competibacter-associated *prpE*, suggesting an advantage for Accumulibacter to compete for propionate over GAOs as electron donor. This is in accordance with previous studies based on process performance monitoring that suggest that propionate favors PAOs over GAOs (Oehmen et al., 2005). The potentially higher fitness of Accumulibacter over Competibacter in utilizing propionate may explain the higher P-removal per C-uptake as well as the better P-removal efficiency when propionate was applied as the primary carbon source (Table 1). Indeed, the presence of an active GAO1 population did not outcompete the Accumulibacter populations in the utilization of either acetate or propionate; based on *ackA, acs,* and *prpE* gene transcripts, the three Accumulibacter populations accounted for 80-90% gene expression related to activation of these primary carbon sources.

As the utilization of the primary carbon sources was limited for the non-PAO and non-GAO genomes based on acetate and propionate activation gene presence and expression levels (>95% of which mapped to Accumulibacter and Competibacter MAGs), the breakdown of EPS produced by Accumulibacter and endogenous metabolism may support their growth and replication. Utilization of the EPS to support the growth of the flanking heterotrophic populations has also been hypothesized in anammox granules (Lawson et al., 2017). Based on genomic content and patterns of gene expression, we hypothesize that the two highly active non-PAO and non-GAO flanking populations, CH7 and PR6, may use extracellular substances including carbohydrates in EPS excreted by Accumulibacter populations, extracellular proteins, and extracellular serine as their primary carbon sources. Amino acids and cofactors cross-feeding may also occur between the Accumulibacter populations and the two flanking populations of CH7 and PR6 (see Figure S5). CH7 affiliates with the Chloroflexota phylum (Chloroflexi phylum in NCBI taxonomy). Members of this phylum have previously been suggested to be active as scavengers of organic compounds in other engineered systems, including anammox reactors (Kindaichi et al., 2012) and activated sludge (Kragelund et al., 2007).

In addition to their putative role in consuming endogenous organic carbon, metatranscriptomic data suggests that the flanking non-PAO PR6 (Gammaproteobacteria) may also play an important role in aggregate (granule) formation. Granules were formed in this DPAO consortia without intentional selection. Type IV pili are recognized as an important cell surface structure that mediate or regulate the bioprocesses involved in establishing/maintaining the biofilms and microbial aggregates (Maldarelli et al., 2016). Genes *PilQ/PilY1* involved in Type IV pilus synthesis were among the most highly expressed genes in PR6. 33% of the mRNA reads mapped to *PilQ/PilY1* associated with PR6, and this ratio was even higher than those in the range of 14%-27% *PilQ/PilY1* mRNA transcripts associated with the three *Accumulibacter* PAO populations. Consistent with the hypothesis that PR6 may support granule formation or structure, genus *Pseudoxanthomonas,* to which PR6 affiliates, has previously been identified as highly abundant in aerobic granular sludge (Weissbrodt et al., 2012; Leventhal et al., 2018).

## Materials and Methods

### Bioreactor operation

A 14L lab-scale denitrifying Enhanced Biological Phosphorus Removal (EBPR) sequencing batch reactor with 12 L working volume was operated continuously with synthetic municipal wastewater for 3 years to select for high denitrifying PAO (DPAO) activity (Gao et al., 2017). The reactor was inoculated with activated sludge from an EBPR process at the Stickney Water Reclamation Plant (Chicago, IL), and operated with an HRT of 12 hours under cyclic anaerobic and anoxic conditions with a short aerobic polishing step. No biomass was intentionally wasted during the operation. Briefly, during the anaerobic period (90 min), the primary carbon source was switched between acetate and propionate (120-150 mg-COD/L) every two SBR cycles. In the subsequent anoxic phase (150 min), high NO_2_ feed (40-50 mg-N/L) intended to mimic effluent from an upstream nitritation reactor was dosed as the terminal electron acceptor for denitrifying phosphate uptake. A 60-min aerobic polishing period was added after the anoxic phase to improve P removal. The initial P concentration was between 5-10 mg PO_4_ -P/L. Detailed reactor operation conditions and performance were described previously (Gao et al., 2017). While the SBR was operated for retention of suspended growth (floccular) biomass, over the course of long-term reactor operation granule formation was observed. During the period of biomass collection for this study, both granules and floccular biomass were present in the reactor.

### Sample collection

To profile gene expression in typical denitrifying EBPR cycles using either acetate or propionate as the primary carbon source, biomass samples were collected every 15 min over complete SBR cycles for RNA extraction and key metabolites profiling. For each carbon source, sampling was conducted in duplicate cycles. Bulk biomass (2 ml) was collected for each time point in microcentrifuge tubes, and preserved using RNA*later* Stabilization Solution (Invitrogen, USA). Samples were centrifuged, supernatant removed, and the pellet submerged in 1 ml of RNA*later* solution overnight. RNA*later* solution was then removed and cell pellets were flash frozen in liquid nitrogen and stored in -80°C freezer prior to RNA extraction. To recover metagenome assembled genomes (for mapping of metatranscriptome sequencing data) and facilitate evaluation of population segregation between aggregate size classes, three biomass samples including one for the bulk biomass and two samples representing different biomass size fractions were collected from the reactor for DNA extraction during the period when gene expression studies were conducted. Because granulation was observed during the long-term reactor operation, aggregate size fraction cutoffs were chosen based on particle size analysis of the biomass (Figure S8). Biomass was then separated based on particle size (<350µm, corresponding to approximately 50% of the total volume, and ≥350µm) by sieving, as described previously ≥ (Gao et al., 2019). Each biomass sample was centrifuged to remove supernatant and stored at -80°C prior to DNA extraction.

### Chemical analyses

Chemical analyses were performed at the same time points used for RNA sample archiving. To monitor the EBPR cycle, NO_2_ and phosphate (PO_4_) were measured with an automated Continuous Flow Analyzer (CFA) (Skalar, Netherlands) based on standard colorimetric methods (APHA, 1998). N_2_O was monitored using an N_2_O-Wastewater sensor (Unisense, Denmark) continuously during the operation. Acetate and propionate were measured using a GC-FID (Thermo Fisher Scientific, USA) following previously described methods (Gao et al., 2017). MLSS and MLVSS were measured using standard methods (APHA, 1998).

### Nucleic Acid Extraction and Metagenomic and Metatranscriptomic Sequencing

Genomic DNA was extracted from each biomass sample in duplicate using the FastDNA SPIN Kit for soil (MP Biomedicals, USA), according to the manufacturer’s protocols. Total RNA was extracted from each time point using the MagAttract PowerMicrobiome RNA kit (Qiagen, USA). Extracted RNA was quality-checked using agarose gel electrophoresis and Synergy HTX Multi-Mode Reader (BioTek, USA). Samples were selected for metatranscriptomic sequencing from each phase of operation (anaerobic: 30 minutes, anoxic: 150 minutes, aerobic: 270 minutes) based on RT-qPCR results of *nirS* and *nosZ* in 15 minute increments (see Table S2 in the supporting information for details). The stranded DNA-seq and mRNA-seq was conducted in the Northwestern University NUSeq Core Facility. Briefly, the DNA and RNA concentrations were determined with a Qubit fluorometer. Total RNA samples were also checked for fragment sizing using an Agilent Bioanalyzer 2100. The Illumina TruSeq Stranded Total RNA Library Preparation Kit was used to prepare sequencing libraries from 1 ug of total RNA samples. This procedure includes rRNA depletion with Ribo-Zero Bacterial rRNA Removal Kit, cDNA synthesis, 3’ end adenylation, adapter ligation, library PCR amplification and validation. An Illumina HiSeq 4000 Sequencer was used to generate paired-end 150 bp reads from the libraries, with an average insert size of 650 bp. Raw paired-end reads were initially filtered, and adapters trimmed using cutadapt v1.13 to remove low-quality bases and adapters from both ends and to discard reads based on maximum error rate of 0.1 and minimum length of 20 bp (Martin, 2011).

### Metagenomic Assembly, Genome Binning, Annotation, and Metabolic Reconstruction

Clean DNA paired-end reads from each sample were assembled separately as well as co-assembled using the *de novo* assembler IDBA-UD using multiple kmer values from 20 to 80 and minimal contig length of 500 bp (Peng et al., 2012). The quality of assembled contigs with different kmers was checked by QUAST (Gurevich et al., 2013). Co-assembled contigs resulted in a maximum N50 and were therefore used for downstream analysis. Coverage across all contigs was calculated by mapping raw reads from each sample against the assembled contigs using Bowtie2 with default parameters (Langmead and Salzberg, 2012). As is summarized in Table S3, shotgun metagenomic sequencing all three samples generated a total of 51.6 Gbp of raw sequencing reads and 49.4 Gbp of clean data after quality-filtering. Co-assembling all the samples yielded a total of 290,166 contigs with an N50 of 4,145. On average, 82.6±1.3% of the clean reads from each DNA sequencing sample and 75.2±1.0% of the filtered mRNA sequencing samples were aligned to the co-assembled contigs, indicating that the co-assembled contigs captured the majority of the metagenomic and metatranscriptomic sequencing reads. Open reading frame (ORF) calling was performed using Prodigal v2.6.2 (Hyatt et al., 2012). To extract draft metagenome assembled genomes from the co-assembled contigs, genome binning was then performed with MetaBAT under the ‘superspecific’ mode with the resulting mapping files, which yielded 32 MAGs. The quality of each draft genome was checked using CheckM v1.0.7 based on 111 essential single-copy genes (Parks et al., 2015). High quality draft genomes with completeness greater than 85% and contamination less than 5% were retained for downstream analysis. Draft genome annotation was performed using Prokka v1.12 (Seemann, 2014) and RAST v0.0.12 (Meyer et al., 2008) in KBase. Genes involved in key metabolic pathways of interest including carbon, nitrogen, phosphorus metabolism, and electron transport were validated manually with the assistance of the BioCyc pathway database and KEGG pathway (Kanehisa and Goto, 2000; Caspi et al., 2012).

### Phylogenetic Analysis of Metagenomic Assembled Genomes

The taxonomic affiliation of each MAG was inferred using GTDB-Tk v0.1.3 under ‘classification’ mode, based on 120 ubiquitous single-copy protein sequences (Parks et al., 2018). To generate a maximum likelihood tree, marker genes from 251 genomes that are closely related to the MAGs in the GTDB reference database were extracted. MAFFT was used to realign the concatenated single-copy protein sequences, and RAxML v8.2.10 was then applied to generate a maximum likelihood tree with the automatic protein model assignment algorithm (PROTGAMMAAUTO) and 100 rapid bootstraps (Stamatakis, 2014).

### ppk1 clone library construction and sequencing

To characterize clade-level community structure of Accumlibacter, a 1123 bp fragment of the *Accumulibacter*-specific *ppk1* gene was PCR amplified from the DNA extracts of bulk biomass using primers ACC-ppk1-254f and ACC-ppk1-1376r (McMahon et al., 2007), as previously described (Gao et al., 2017). Triplicate PCR products were pooled and purified via gel electrophoresis using the PureLink Quick Gel Extraction Kit (ThermoFisher). The purified PCR products were cloned using the TOPO® TA cloning® kits (Invitrogen), as per the manufacturer’s protocol. Among the white colonies produced, 35 colonies were picked and cultured overnight in LB medium containing 50 μg/ml ampicillin. The plasmids were isolated using the PureLink Quick Plasmid Miniprep Kit (ThermoFisher) and sequenced by ATGC Inc. (Wheeling, IL) with an ABI 3730 DNA analyzer (Applied Biosystems, USA).

### ppk1 gene screening and phylogenetic analysis

To identify *ppk1* genes in assembled metagenomic contigs, a BLAST+ search was performed using ORFs predicted from co-assembled contigs as a query, against 781 reference Accumulibacter-associated *ppk1* sequences based on an e-value cut-off of 10^-5^ (Camacho et al., 2009).13 unique *ppk1* genes were annotated from the metagenomic dataset. The micro-diversity of these 13 assembled *ppk1* genes and the 35 *ppk1* gene clones were evaluated via phylogenetic analysis. 952 *ppk1* gene sequences downloaded from NCBI were applied as reference *ppk1* sequences. The *ppk1* genes were aligned through MAFFT (Katoh and Standley, 2013) over a consensus region of 1007 bp. Maximum-likelihood phylogenetic trees were constructed with RAxML v8.2.10 with the automatic protein model assignment algorithm (PROTGAMMAAUTO) and 100 rapid bootstraps (Stamatakis, 2014). Visualization of the phylogenetic tree was through the iTOL platform (Letunic and Bork, 2016).

### Metatranscriptomic analysis

Ribosomal rRNA sequences in the clean metatranscriptomic reads were identified by aligning the clean reads against the SILVA (SSURef and LSURef) database release 128 using BLAST+ (Quast et al., 2013; Yilmaz et al., 2014). Sequences with alignments to the rRNA database in SILVA that had an e-value < 1e-9 were assumed to be rRNA and discarded from further analyses. The resulting non-rRNA (mRNA) reads were mapped to all co-assembled metagenomic contigs using Bowtie2 with the end-by-end mode (Langmead and Salzberg, 2012). To increase the mapping sensitivity, the following parameters were used when applying Bowtie2: the length of the seed substrings to align=25, maximum mismatch penalties=10, and minimum mismatch penalties=5.

To profile gene expression across the MAGs, htseq-count v0.6.1 was used to calculate the mRNA read counts mapped to each predicted ORF with the ‘intersection strict’ parameter (Anders et al., 2015). mRNA-RPKM values of each ORF were then calculated by normalizing the mapped mRNA read counts by the sequencing depth and ORF length. Pathway expression levels were calculated based on averaging the RPKM values for each gene involved. To compare gene expression dynamics under different redox conditions (anaerobic/anoxic/aerobic), the gene expression change for metabolic pathways or functional genes was determined by normalizing the RPKM values under the anoxic or aerobic conditions to the RPKM values under the anaerobic conditions (mRNA/mRNA ratio).

### Carbohydrate hydrolase and membrane transport proteins identification

Carbohydrate hydrolases and membrane transport proteins were identified in each genome by BLASTP searches against the CAZy (Lombard et al., 2014) and TCDB (Saier et al., 2016) databases. CELLO v2.5 was used to predict the subcellular location of identified carbohydrate hydrolases using support vector machines based on n-peptide composition (Yu et al., 2004).

### Data availability

Raw metagenome DNA (3 datasets in total) and metatranscriptome RNA (6 datasets in total) sequencing data are available in the NCBI Sequence Read Achieve (SRA) database under the Bioproject accession number of PRJNA576469. GeneBank accession numbers of the 35 *ppk1* gene clone sequences are MN551751-MN551785. Accession numbers of the 32 MAGs recovered in this study are SAMN12995720-SAMN12995751.

