## supporting information for "Integrated Omics Analyses Reveal Differential Gene Expression and Potential for Cooperation Between Denitrifying Polyphosphate and Glycogen Accumulating Organisms"

**Table S1.** ANI between the draft metagenome assembled genomes from this study and the corresponding nearest publicly available reference genome (NCBI accession numbers provided in the table).

| **Bin_ID** | **Reference genome** | **ANI** |
| --- | --- | --- |
| Acc-IF | UBA11064 | 83 |
| GAO1 | GCA_002391525.1 | 78 |
| PR6 | GCF_000972865.1 | 87 |
| CH7 | GCA_002746795.1 | 77 |
| BA1 | GCA_001897835.1 | 76 |
| CH1 | GCA_001567085.1 | 76 |
| CH6 | GCA_002699125.1 | 76 |
| CH2 | GCA_002352035.1 | 76 |
| BA12 | GCF_002283555.1 | 76 |
| BA2 | GCA_002352145.1 | 76 |
| VE2 | GCA_002396485.1 | 76 |
| CH3 | GCA_002842085.1 | 76 |
| MY1 | GCF_000331735.1 | 77 |
| VE1 | GCA_002396485.1 | 77 |
| VE3 | GCF_000972765.1 | 77 |
| PL1 | GCA_002343325.1 | 78 |
| BA6 | GCA_002344125.1 | 78 |
| CH5 | UBA9854 | 79 |
| PR2 | GCA_002280405.1 | 79 |
| PR4 | GCF_001293525.1 | 79 |
| PR3 | GCA_001770955.1 | 79 |
| CH4 | GCA_001567085.1 | 79 |
| VE4 | GCA_002344865.1 | 80 |
| PR5 | GCF_900155935.1 | 84 |
| BA10 | GCA_002839825.1 | 86 |
| BA7 | UBA11072 | 96 |
| BA8 | GCA_002344125.1 | 98 |
| BA3 | GCA_002344975.1 | 98 |
| BA5 | GCA_002426785.1 | 98 |
| PR1 | GCA_002425415.1 | 99 |
| BA11 | GCA_001567175.1 | 100 |

**Table S2.** RT-qPCR quantification on the expression of the *nirS* and *nosZ* gene during typical SBR batches with either acetate or propionate as the primary carbon source. Fold change is relative to the 5 minute time point during the anaerobic period. Based on RT-qPCR results shown here, time points 30 minutes (anaerobic), 150 minutes (anoxic), and 270 minutes (aerobic) were selected for metatranscriptomic sequencing.

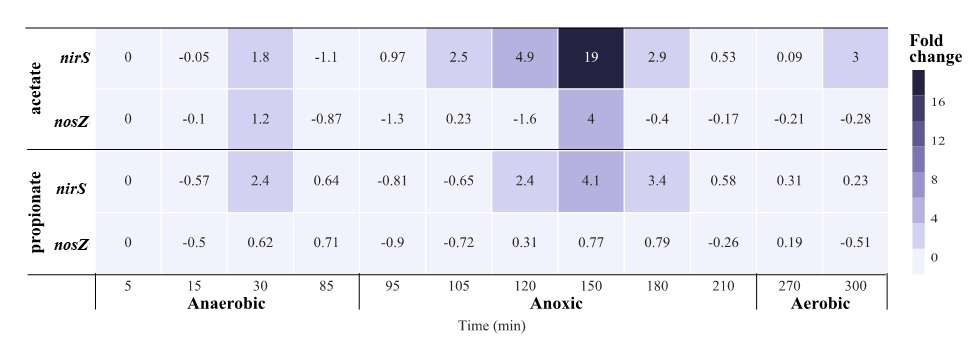

**Table S3.** Summary statistics for metagenomic sequencing and assembly

|  | **Total reads (Gbp)** | **Filtered Reads (Gbp)** | **kmer** | **Total assembled reads (Gbp)** | **Longest contig** | **# of Contigs** | **N50* (Contigs)** |
| --- | --- | --- | --- | --- | --- | --- | --- |
| DNA-Total^*^ | 9.1 | 8.6 | 70 | 0.36 | 331,655 | 165,316 | 3,707 |
| DNA-L-350 | 27.1 | 26.3 | 70 | 0.26 | 209,078 | 129,208 | 2,905 |
| DNA-S-350 | 15.5 | 14.5 | 70 | 0.43 | 331,566 | 174,811 | 4,643 |
| co-assembly | 51.7 | 49.4 | 70 | 0.66 | 412,072 | 290,166 | 4,145 |

^*^DNA-Total: Bulk sludge DNA extracts

DNA-L-350: DNA extracts subjected to the metagenome sequencing from sludge particulates with diameter > 350 µm

DNA-S-350: DNA extracts subjected to the metagenome sequencing from sludge particulates with diameter < 350 µM.

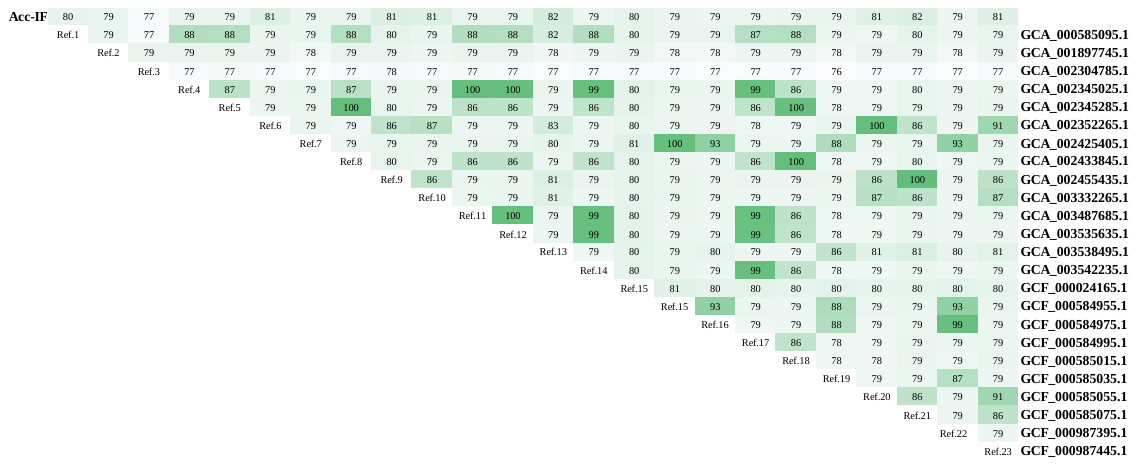

**Figure S2.** ANI (%) between Acc-IF and the 24 publicly available Accumulibacter (PAO) reference genomes (accession number shown on the far right)

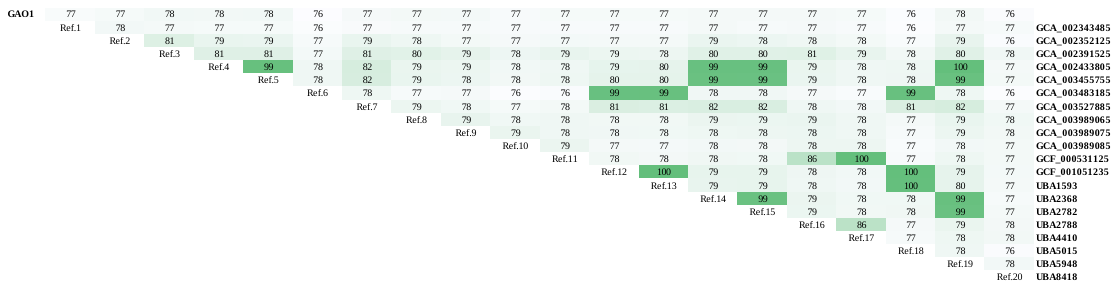

**Figure S3.** ANI (%) between GAO1 and the 19 publicly available Competibacter (GAO) reference genomes (accession number shown on the far right)

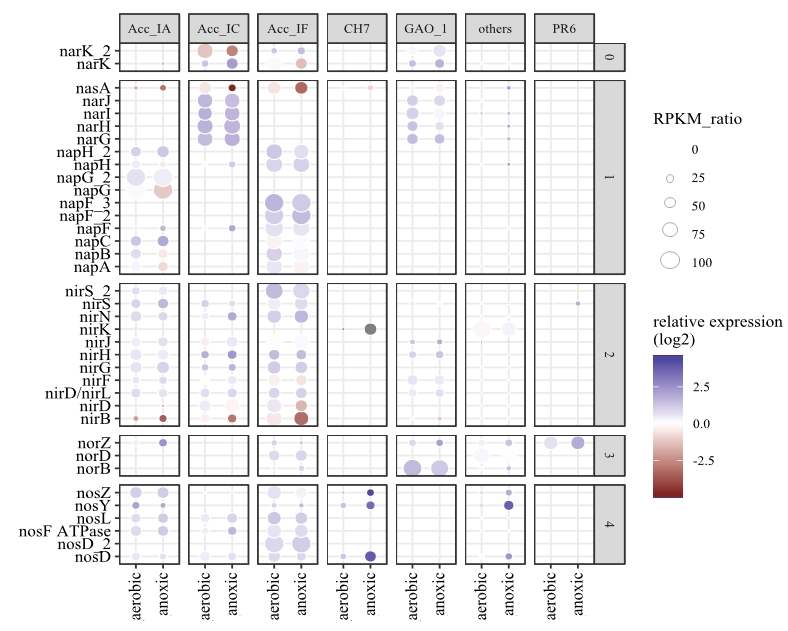

**Figure S4.** Denitrification gene expression under different redox conditions. The relative gene expression under anoxic and aerobic conditions was normalized to the RNA-RPKM value under anaerobic conditions. The relative gene expression is averaged across propionate and acetate fed cycles. Blue indicates up-regulation and red indicates down-regulation, relative to anaerobic conditions. Abundances of the gene transcripts expressed by each MAG are indicated by the size of the bubbles. 0- Nitrate/Nitrite transporter, 1- Nitrate reduction, 2- Nitrite reduction, 3- Nitric oxide reduction, and 4- Nitrous oxide reduction.

Ratio of each clade in sludge particulates (%)

| ppk1 gene clades identified in this DPAO consortia | Size <350µm | Size>=350µm |
| --- | --- | --- |
| IA | 17.5 | 37.7 |
| IB | 8.4 | 5.7 |
| IC | 36.1 | 25.6 |
| ID | 7.4 | 13.9 |
| IF | 30.6 | 17.1 |

Figure S1. Distribution of the *Accumulibacter* clades based on *ppk1* gene phylogenetic analyses in larger sludge particulates (granules with diameter >=350 µm) and in smaller sludge particulates (diameter < 350 µm)

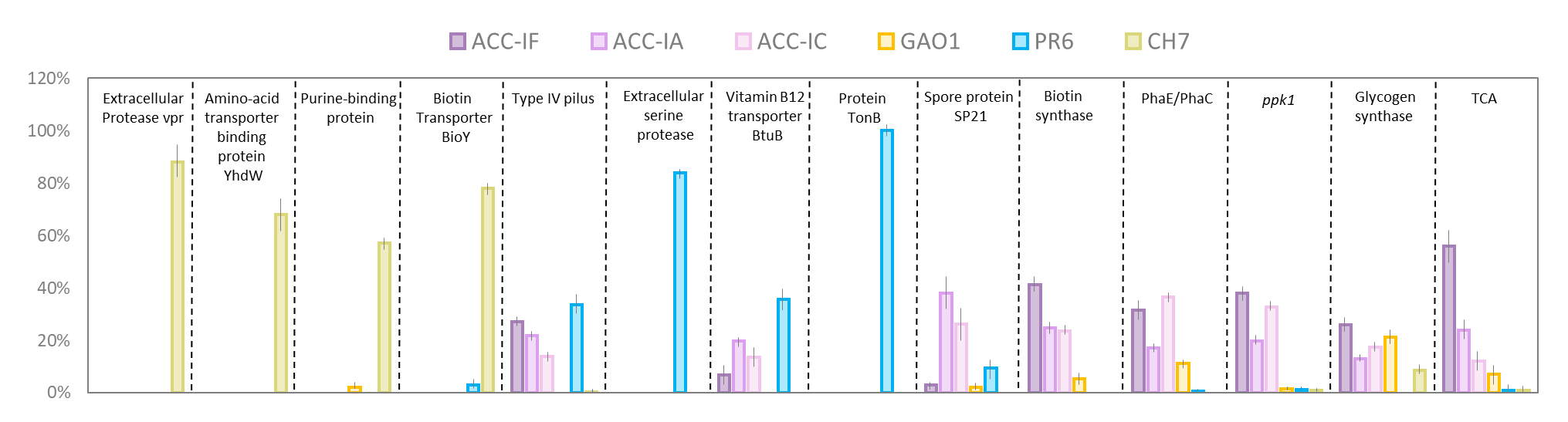

**Figure S5.** RNA-RPKM based quantification of gene transcripts associated with the key metabolism in each of the 6 identified highly active MAGs. Gene transcripts expressed by different MAGs were presented in different colors, and the height of each bar represents the proportion of gene transcripts expressed by each MAG. The RNA-RPKM values were averaged across the six metatranscriptome datasets and the specific RNA-RPKM values of genes associated with the metabolism in this figure could be found in the appendix file 1.

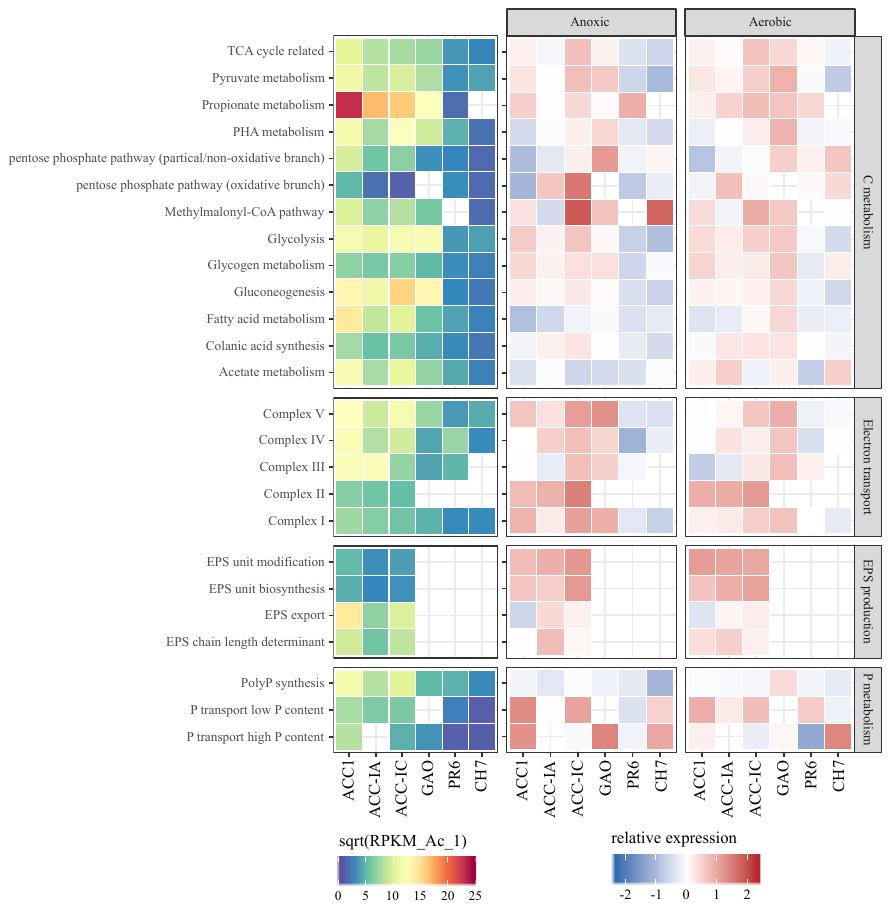

**Figure S6.** Gene expression patterns (RPKM and relative) of major carbon, electron transport, EPS production and P metabolic pathways by each dominant MAG. The 1^st^ column on the left represents the RNA-RPKM values of the genes involved under the anaerobic period, and colors changing from purple to red represents the increase of RPKM values. The 2^nd^ and 3^rd^ columns represent relative gene expression under the anoxic and aerobic conditions, normalized by the RNA-RPKM values under the anaerobic condition. MAGs with missing genes in the pathways are marked as blank in the heatmap.

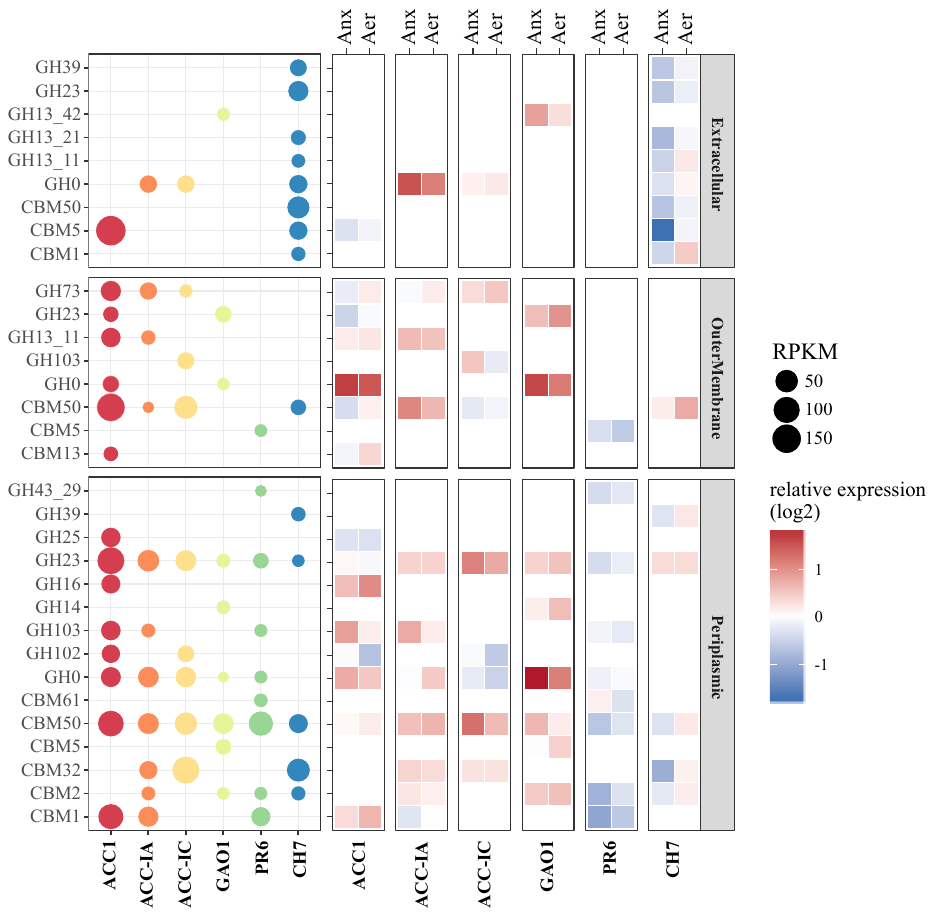

**Figure S7.** CAZy gene expression in the 6 dominant MAGs (in terms of gene expression). The 1^st^ column on the left summarizes the RNA-RPKM values of each gene under the anaerobic period, where the size of each bubble represents the RNA-RPKM values. The 2^nd^ and 3^rd^ columns represent relative gene expression under the anoxic and aerobic conditions, normalized by the RNA-RPKM values under the anaerobic condition. GH: glycoside hydrolases, CBM: carbohydrate-binding module

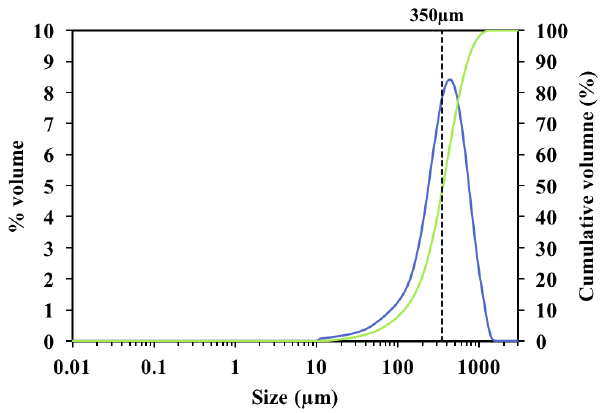

**Figure S8.** Distribution of the aggregate size characterized via particle size analyzer. Blue indicate volume %, and green indicates cumulative volume %.
